# Transcriptomic changes due to early, chronic alcohol exposure during cortical development implicate regionalization, cell-type specification, synaptogenesis and WNT signaling as primary determinants of fetal alcohol Spectrum Disorders

**DOI:** 10.1101/784793

**Authors:** Máté Fischer, Praveen Chander, Huining Kang, Jason P. Weick

**Affiliations:** Department of Neurosciences, University of New Mexico HSC, Albuquerque, NM 87131, USA; Center for Brain Recovery and Repair, University of New Mexico HSC, Albuquerque, NM 87131, USA

**Keywords:** Regionalization, patterning, WNT signaling, FASD, hPSN

## Abstract

Fetal alcohol spectrum disorders (FASD) are described by a cluster of deficits following *in utero* alcohol exposure, whose effects disproportionately target the cerebral cortex. *In vitro* and *in vivo* models of FASD have successfully recapitulated multiple facets of clinical presentations, including morphological and behavioral deficits, but far less is understood regarding the molecular and genetic bases of FASD. In this study, we utilize an *in vitro* human pluripotent stem cell-based (hPSC) model of corticogenesis to probe the effect of early, chronic alcohol exposure on the transcriptome of developing cortical neurons. We here identify a relatively limited number of significantly altered biological pathways, including regional patterning, cell-type specification, axon guidance and synaptic function. Significant upregulation of WNT signaling-related transcripts, to the exclusion of other secreted morphogens was also observed in alcohol exposed cultures. Lastly, an overall alcohol-associated shift towards an increased caudal profile, at the expense of rostral molecular identity was observed, representing a potentially previously underappreciated FASD phenotype.

## Introduction

Fetal alcohol spectrum disorders (FASD) refer to a cluster of physical and mental symptoms affecting a person exposed to alcohol during gestation. FASD is an umbrella term that encompasses fetal alcohol syndrome (FAS), partial FAS, alcohol related neurodevelopmental disorder (ARND) and alcohol related birth defects (ARBD) (Kodituwakku 2007). According to estimates from the CDC, accidental and intentional drinking during pregnancy affects approximately 1.5% of the world population, and the added cost of care for individuals with FASD in the U.S. is estimated at $2 million over the course of their lifetime (Popova 2017). FASD can present with a large variety of severity of symptoms, from relatively mild perturbations to adaptive learning, attention, executive function, social cognition and craniofacial morphogenesis, to severely debilitating disabilities (Lange 2017). Abstinence from alcohol during pregnancy can completely prevent the development of the disorder, but as many women do not discover they are pregnant for several weeks after conception, it is critical to understand the types of neurological insults that can occur before that time (CDC 2009).

The onset of neurogenesis in the cerebral cortex occurs during the first trimester in humans and injury during this period can result in various cortical malformations. The cerebral cortex is thought to be the major target of prenatal alcohol exposure (PAE), as many of the social, affective and cognitive deficits exhibited by children with FASD are mediated by cortical regions. Various research models of PAE have confirmed that alcohol produces significant alterations in cortical development at the gross morphological, cellular, and subcellular levels (Granato 2018). Impaired proliferation of radial glial cells in response to alcohol exposure has been shown to underlie a reduction of neurogenesis at high doses (70-100 mM), but appears to be less affected at doses ≤50mM (Zecevic 2012; Larsen 2016). Human pluripotent stem cell-derived neurons (hPSN) have been demonstrated to recapitulate multiple aspects of *in utero* neuronal patterning and specification, making them an ideal model system for investigation of such early developmental questions (Erceg 2009, Zhang 2010, Studer 2017). Critically, default differentiation without the addition of patterning morphogens produces cerebral cortical neurons on a physiologically-relevant time scale (Vanderhaeghen 2015). Similar, more targeted studies have previous come to a wide variety of conclusions concerning the effects of alcohol on proliferation, cell type specification, as well as synaptogenesis, although depending on the cell lines and dosing paradigm employed, these results can often be contradictory (Yang 2012, Leigland 2013, Treit 2014).

Using qPCR-based RNA analysis, our lab has previously demonstrated that chronic 50mM alcohol exposure to developing hPSNs leads to significant alterations in a number of mRNAs associated with ventral forebrain patterning and GABAergic neuron specification (Larsen 2016). However, this was not accompanied by overall changes to the number of GABAergic interneurons generated, nor was a functional excitatory-inhibitory (E/I) imbalance uncovered. Thus, to probe for potential compensatory mechanisms and/or other mechanistic underpinnings of FASD-associated neurodevelopmental deficits, we used bulk RNA-sequencing to determine how alcohol affects RNA expression at the transcriptome level. These current data support previous findings concerning targeted downregulation of transcripts related to GABAergic patterning and excitatory/inhibitory balance. Additionally, we report significant perturbation to transcripts associated with WNT signaling and cortical regionalization, leading to an overall more caudal forebrain signature with alcohol exposure.

## Results

### Differentiation of Human Neurons and Global Transcriptomic Findings

Figure 1A demonstrates the developmental timeline of neuronal differentiation from hPSCs to functional cortical neurons. To generate mixed cortical cultures we employed a modified default differentiation paradigm using the serum-free embryoid body (SFEB) that results in an >95% Pax6^**+**^ population of forebrain neuroectoderm following 10 days of differentiation, and functional post-mitotic neurons by 7 weeks of differentiation similar to *in vivo* human cortex (Zhang 2001, Lavaute 2009, Weick 2011 and 2016). Without the application of exogenous morphogens the H9 cell line produces cortical a mixed culture comprised of glutamatergic projection and GABAergic interneurons as well as neural progenitor cells, all with a primarily cortical pattern of gene expression (Floruta 2017, Nadadhur 2018). To determine the effect of alcohol on neuronal specification and patterning we applied 50mM alcohol daily throughout the differentiation protocol similar to previous reports (Larson 2016), which was meant to mimic an early exposure during the periods of gastrulation and neurulation similar to *in utero* first-trimester chronic binge exposures (Okada 2009, Vaccarino 2012).

**Figure 1.**
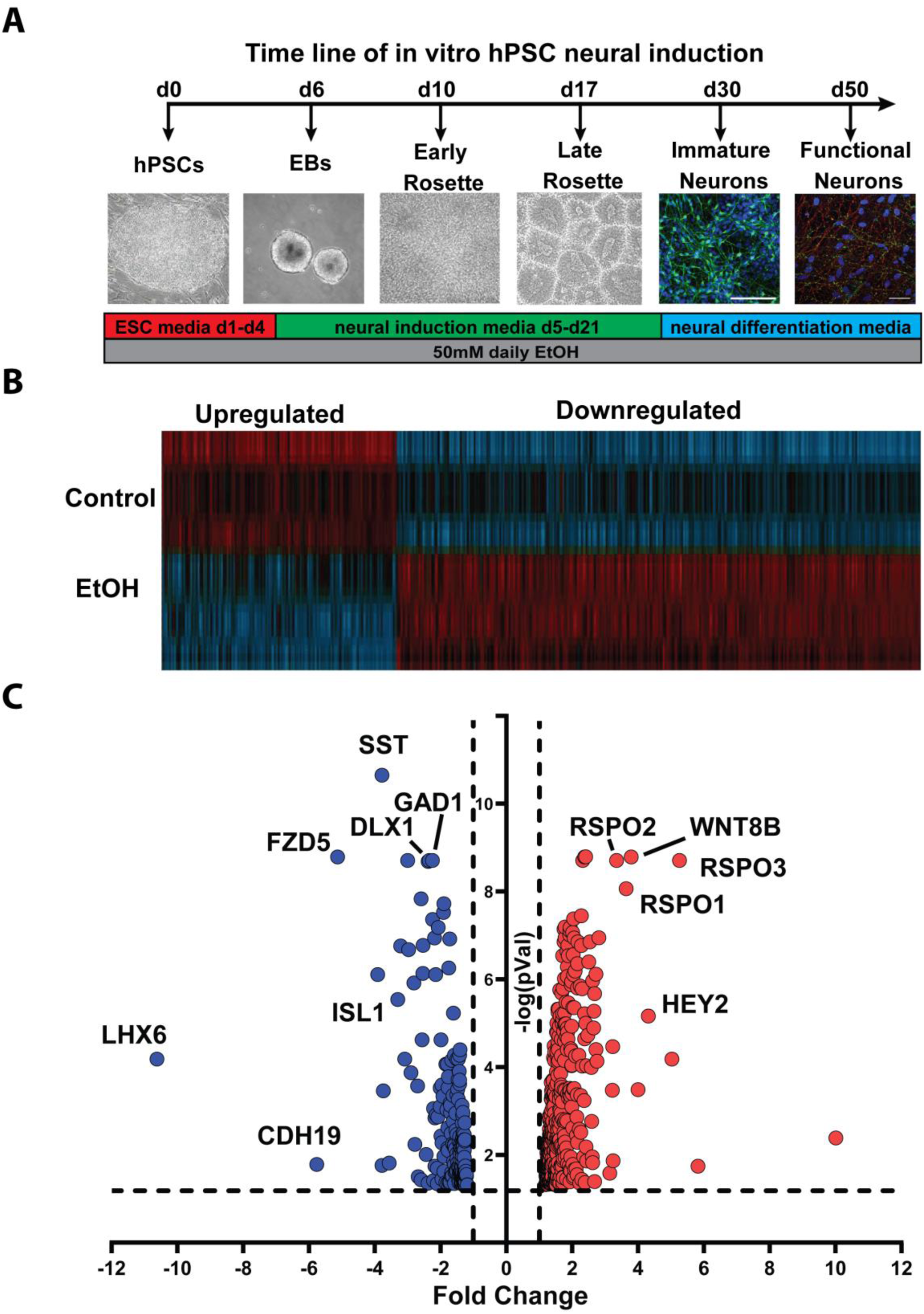

Gene-level data was compared for three biological replicates per treatment group and a DE gene list was assembled based on cutoff values for 1.2-fold change and statistical significance (adjusted p-value<0.05). After these filters were applied, 691 mRNA transcripts were revealed to be significantly altered between untreated and alcohol-treated cells at day 50 (Fig. 1B). Unbiased hierarchical clustering of DE genes by treatment group (Fig. 1B) demonstrates that alcohol exposure led to robust and highly reproducible patterns of change in the expression of these transcripts across the three replicates. Interestingly, alcohol had an overall positive impact on mRNA expression similar to previous reports (Qin 2017), with 477 transcripts upregulated compared to just 214 downregulated. Splicing analysis was performed for all DE mRNAs, which revealed changes to both major and minor species. While several mRNAs showed significant alternative splicing of major isoforms in complementary directions, the majority were altered uniformly across isoforms (Supplemental Figure 1; Table1). The asymmetrical effect favoring expression increases is illustrated by volcano plot (Fig. 1C), which also highlights the most differentially expressed (DE) gene transcripts, including a dramatic upregulation of WNT co-agonists R-spondin family members (RSPO1-3) as well as WNT8. In contrast, multiple GABAergic interneuron-related transcripts including DLX1, GAD1, and somatostatin (SST) as well as the WNT receptor FZD5 constitute some of the most significantly downregulated transcripts (Fig. 1C).

### Canonical Signaling Pathways Altered with alcohol Exposure

To gain insight into whether alcohol selectively altered the expression of genes with common biological motifs, the list of DE transcripts was first analyzed using DAVID, a NIH-supported suite of bioinformatic tools (v6.8; Huang 2009). Functional annotation of DE transcripts utilizing the gene ontology (GO) algorithm identified significant categories of enrichment (Supplemental Table 2). Some of the most selectively enriched GO categories are listed in Figure 2A and seem to revolve around patterning of the cortical protomap, such as “rostral/caudal axon guidance,” “forebrain rostral/caudal pattern specification” as well as “cerebral cortex regionalization” (Fig. 2A). In addition to regional specifiers, abnormalities in neuronal cell fate decisions were highlighted with the emergence of categories such as “commitment of neuronal cell to specific neuron type in forebrain” and “cerebral cortex GABAergic interneuron differentiation.” Taken together, these findings highlight previous changes observed that point to alterations in excitatory/inhibitory (E/I) cell patterning, but also implicate more global alterations to cortical regionalization.

**Figure 2.**
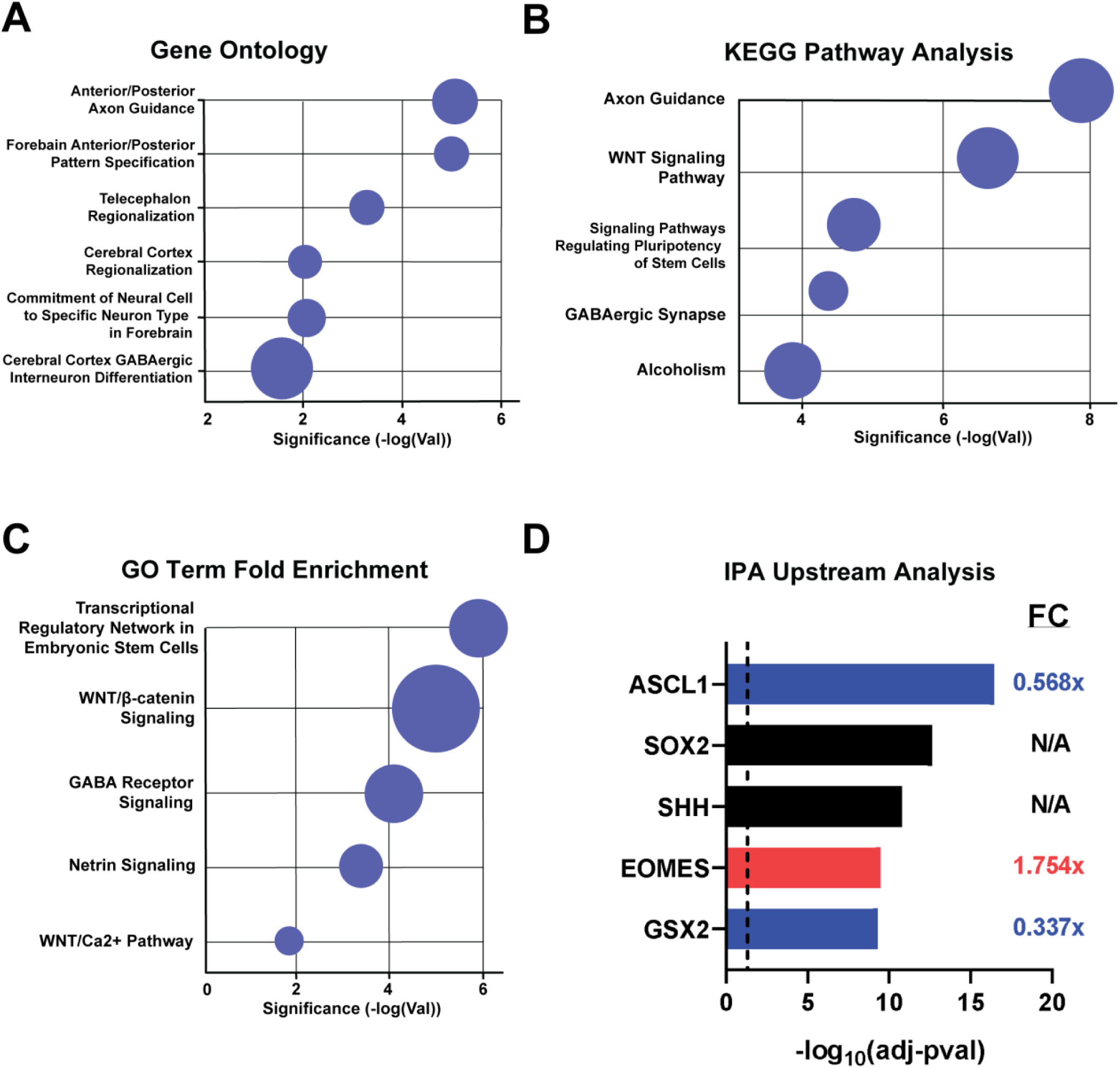

The Kyoto Encyclopedia of Genes and Genomes (KEGG) analysis tool supported by the Kanehisa Laboratory in Japan relies on analysis of pathways rather than ontology terms (Kanehisa 2019). “Axon guidance,” “WNT signaling” and “GABAergic synapse” again feature prominently among the results, providing support for the GO outputs. Additionally, “signaling pathways regulating pluripotency of stem cells” and “alcoholism” were also identified, implicating alcohol exposure-derived dysregulation of stem cell differentiation. Lastly, we took advantage of Ingenuity Pathway Analysis (IPA; Qiagen) to further contextualize the changes reported in the DE gene lists with another orthogonal algorithm. These analyses augment the GO and KEGG findings, as the IPA database is manually curated and based on peer-reviewed literature rather than high-throughput, *in silico* data-mining (Kramer 2014). According to IPA’s analysis of the significantly enriched pathways, the most altered categories included WNT signaling, GABA receptor signaling, synaptogenesis and transcriptional regulation of stem cells (Fig. 2C, Supplemental Table 4). In addition to pathway analyses, IPA enables users to identify potential upstream transcriptional regulators of DE genes as well as whether those transcription factors show up or downregulation in the given dataset. Interestingly, the transcription factors *ASCL1* and GSX2 were not only predicted to be regulating hPSN differentiation but were also shown to be 1.76-fold and 2.97-fold downregulated with alcohol, respectively (Fig. 2D, Supplemental Table 5). Importantly, both these transcription factors are known to be involved in GABAergic interneuron patterning, as well as axonogenesis and synaptic patterning (Mizuguchi 2006, Kessaris 2014, Sun 2016). In contrast to ASCL1 and GSX2, EOMES (TBR2) was found to be upregulated 1.76-fold with alcohol, and has been shown to be a critical regulator of cortical glutamatergic neuron differentiation (Arnold 2008, Sessa 2008). SOX2 and SHH, critical regulators of neural progenitor cells and regional patterning of the cortex, were additionally found to be potential upstream targets of alcohol in our dataset. However, these were not significantly altered in the abundance of their transcripts by alcohol; it is worth noting than neither SHH, nor any of the hedgehog ligands, were found to be expressed to detectable levels as demonstrated previously for default-derived neurons from the H9 cell line (Xu 2010, Floruta 2017, Nadadhur 2018).

### Alcohol Exposure Alters Transcripts associated with both GABAergic and Glutamatergic Neuron Differentiation

Multiple studies have previously implicated developmental alcohol exposure and alterations to the expression of several genes known to affect the patterning, differentiation and migration of GABAergic interneurons, including by our group (Yeh 2008, Vangipuram 2011, Larsen 2016). In the current study, our data suggest multiple levels of regulation at which this GABAergic developmental program is altered. The genes encoding for numerous transcription factors (TFs) associated with interneuron development were significantly *down*regulated, including *ASCL1, GSX2, SIX3* as well as multiple members of the DLX family of TFs (*DLX1/2/5/6)* (Fig. 3A). In addition, alcohol downregulated markers of mature post-mitotic neurons (*NPY* and *SST*) as well as both mRNAs coding for the GABA-synthetic enzymes glutamic acid decarboxylase 65/67 (*GAD1/2*) by more than 2-fold (Fig. 3A). The simultaneous upregulation of mature interneuron subtypes (NDNF and RELN) may suggest compensatory changes that maintain E/I balance in default-generated hPSNs exposed to chronic alcohol (Lake 2016).

**Figure 3.**
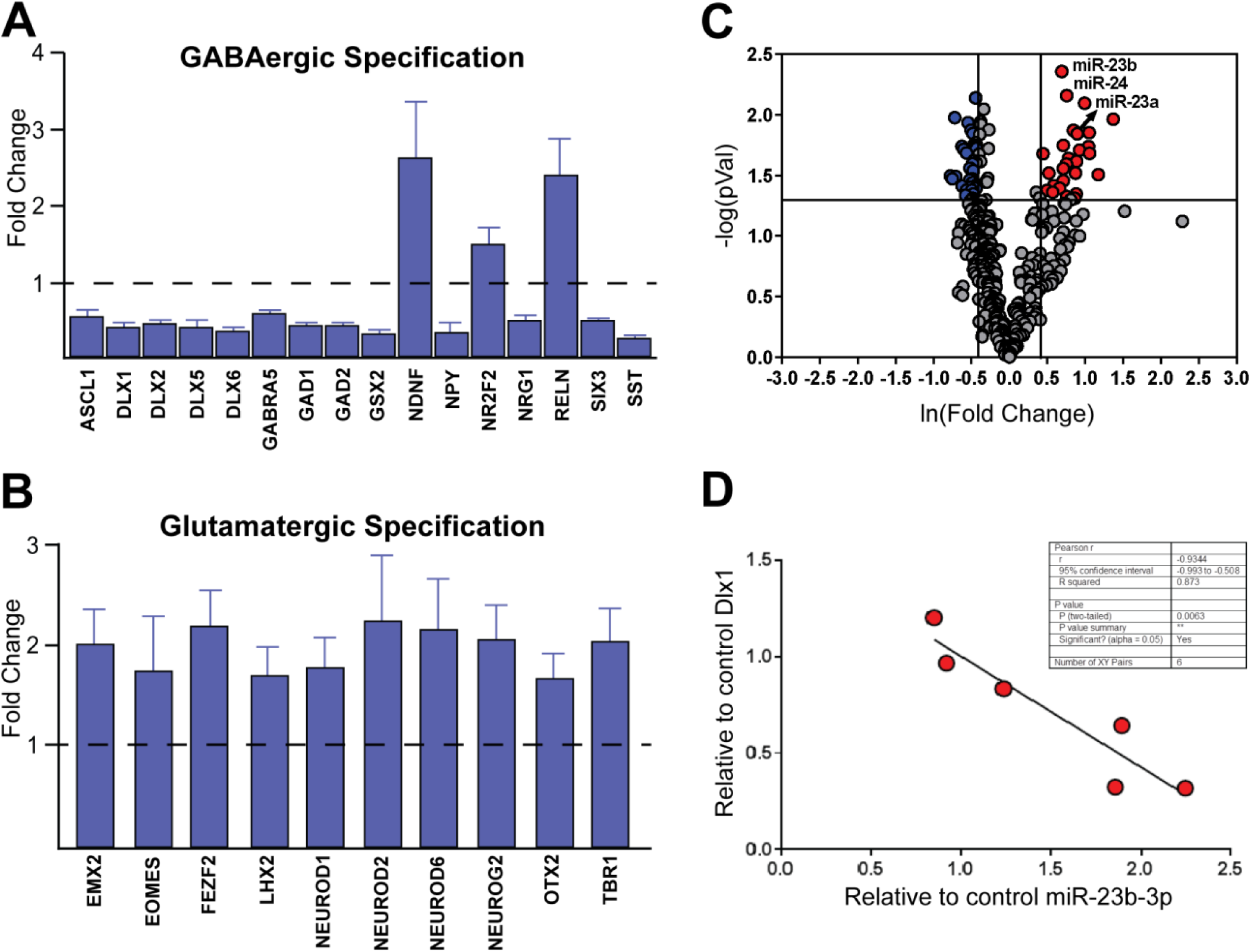

While we did not previously identify glutamatergic patterning genes altered by alcohol using targeted PCR-based analyses (Larsen 2016), other studies have shown that alcohol significantly impacts cortical glutamatergic neuron generation (Qin 2017). Importantly, the current RNA-sequencing analyses did identify a large number of transcripts involved glutamatergic neuron specification to be significantly altered in the alcohol-treated group (Fig. 3B). In contrast to GABAergic transcripts, most glutamatergic mRNAs were upregulated. This included the homeobox domain-containing transcription factors *EMX2, LHX1/2/5/9* and *OTX2*, the zinc finger protein *FEZF2*, basic helix-loop-helix transcription factors *NEUROD1/2/6, NEUROG2* (*NGN2*) and importantly both t-box binding proteins *TBR1* and *TBR2* (*EOMES*). Interestingly, OTX2 as well as the DLX family of TFs, have been recognized for their role not only in brain development, but also craniofacial development, specifically of the rostral aspects of the head (Qui 1997, Wilkie 2001). This is noteworthy as dysregulation of these processes is a hallmark of individuals with FAS (Matsuo 2018). The proteins coded for by a number of these genes have also been identified by other groups as showing altered expression with prenatal alcohol exposure, such as TBR2 (Rakic 2011), FEZF2, NEUROD2, and NEUROG2 (Mandal 2018).

To understand whether microRNAs may play a role in altering mRNA expression we performed nanostring-based sequencing of the same samples that were run for bulk RNA-sequencing. Interestingly, relatively few miRNAs were significantly altered by we discovered a significant two-fold increase in *miR-23b*, a developmentally-regulated miRNA known to be enriched in glutamatergic cells over GABAergic, as *miR-23b* is thought to bind and sequester *DLX1*, repressing its IN-patterning effect (Figure 3C, He 2012). Furthermore, *miR-23b* expression is inversely correlated to *DLX1* expression in these data, suggesting yet another potential level of phenotypic regulation in these developing cortical cells (Figure 3D, Supplemental Figure 2).

### Transcripts involved with synaptic function implicate altered E/I balance in PAE

In addition to the perturbations of genes related to neuronal specification, KEGG and IPA analyses revealed that alcohol had a highly significant effect on mRNAs involved both GABAergic and glutamatergic synaptic function (Figure 2B). Figure 4B shows a diagrammatic summary of data list in Figure 4A. On balance, these data illustrate an overall reduction of gene expression associated with GABA signaling and concurrent increase in glutamatergic transmission-transcripts. Specifically, both enzymes required for the synthesis of GABA (*GAD1* and *GAD2*), along with the vesicular GABA transporter *VGAT*/*SLC32A1*, were all significantly downregulated more than 2-fold. In contrast, both vesicular glutamate transporters (*VGLUT1*/*SLC17A7* and *VGLUT2*/*SLC17A6*) showed concurrent upregulation. Of additional interest were the alcohol-mediated changes to the expression of multiple members of the glypican family (*GPC1, 3, 4*), which are astrocyte-secreted factors that have been shown to be sufficient for inducing functional glutamatergic synapse formation (reviewed by Allen & Eroglu 2017). Collectively, these data add to the hypothesis that altered E/I imbalance may be a crucial mechanism underlying FASD pathologies despite the fact that our previous findings did not reveal gross changes in ratios of excitatory and inhibitory post-synaptic currents (Larsen et al., 2016).

**Figure 4.**
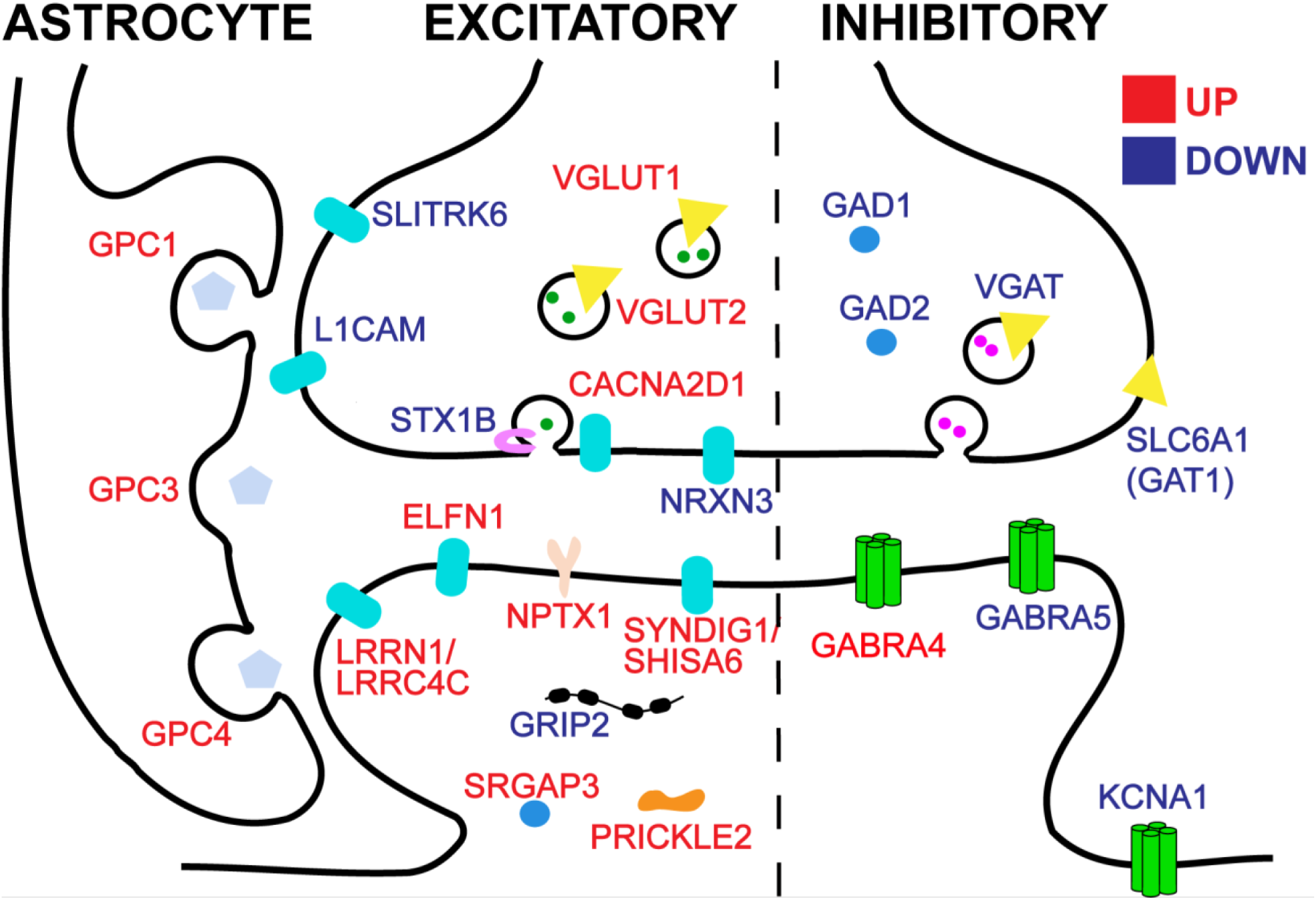

### Alcohol primarily alters the WNT family of Secreted Morphogens

Appropriate spatial and temporal regulation of secreted morphogens is required for proper regionalization, cell-type specification, axon guidance, and synaptic development of the brain and spinal cord during (Wilson 2004, Lumsden 2005). Major morphogen signaling pathways implicated in these processes include the bone morphogenetic proteins (BMPs), fibroblast growth factors (FGF), Notch/Delta, and members of the hedgehog (HH) family (reviewed in Mallamaci 2006). Figure 5A-E highlights selected members of each of these families that have been shown to facilitate cortical patterning and are expressed in default-differentiated hPSN cultures. While some individual transcripts of the BMP (*BMP2/7*), FGF (*FGFR3*), Notch (*NOTCH3*), and HH (*GLI3*) pathways were altered with alcohol, the preponderance of the genes within each pathway remained relatively unchanged with the notable exception of WNT signaling transcripts (Fig. 5E).

**Figure 5.**
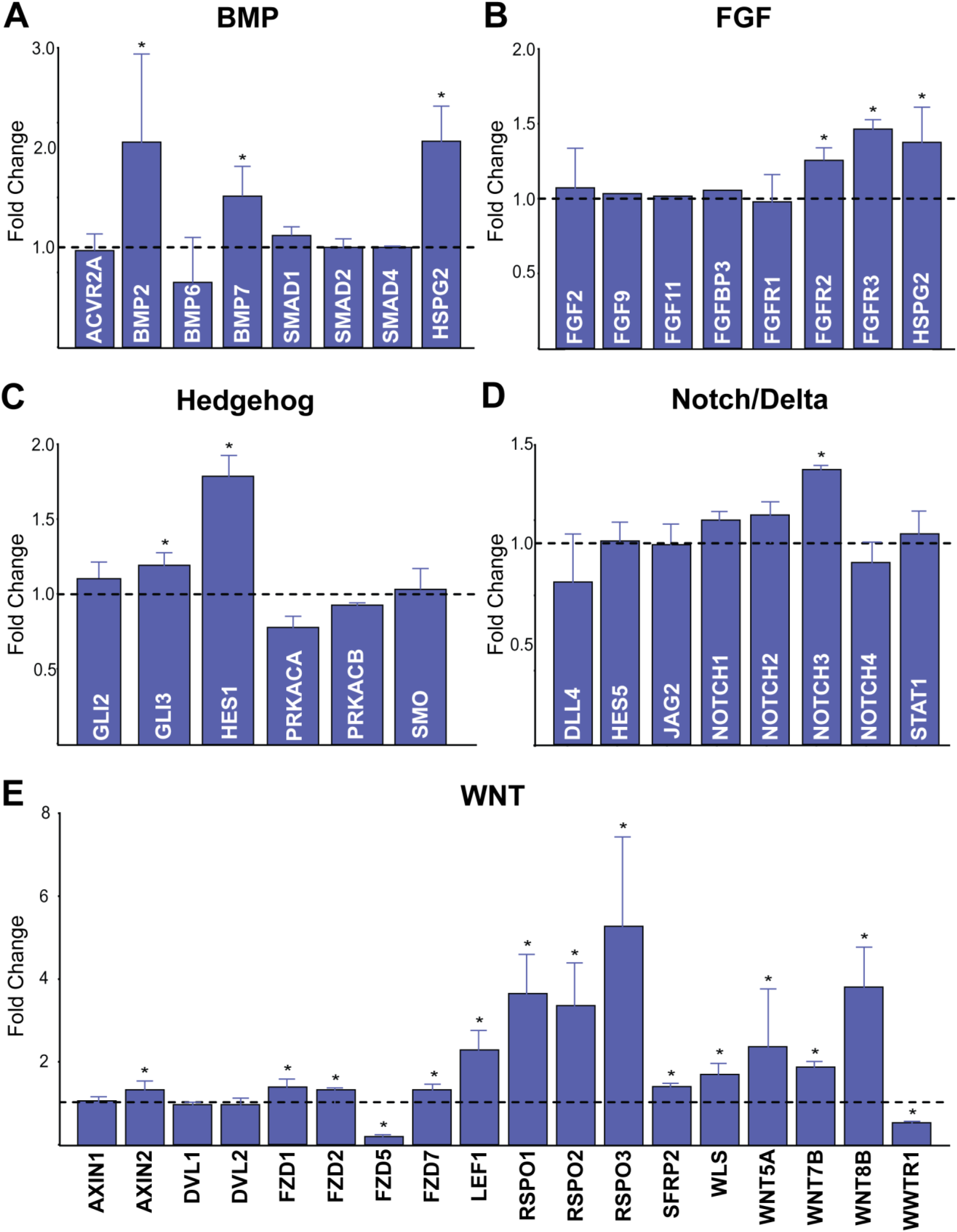

By contrast, transcripts associated with the WNT signaling pathway were identified through unbiased informatic analyses (Fig. 2B, C) as significantly altered in alcohol-treated hPSNs. Overall, fourteen transcripts associated with WNT signaling were altered by alcohol treatment, with 85.7% (12/14) of the genes investigated demonstrating significant upregulation. Interestingly, all three members of the secreted WNT co-agonist R-spondins (*RSPO1-3*) were among the most significantly upregulated transcripts. In addition, the WNT ligand transcripts *WNT5A, 7A and 8B* (Fig. 5E) were also all significantly upregulated. Importantly, not only were secreted ligands altered, downstream transcription factor targets of WNT signaling were also upregulated with alcohol treatment, including *TCF3/4* and *LEF1*. On the other hand, one of the most highly downregulated WNT factors was the Frizzled receptor (*FZD5*), which was lowered to nearly undetectable levels in alcohol-treated cells. However, the more highly expressed *FZD2* receptor showed a minor but significant increase in expression.

### Alcohol exposure specifically increases transcripts with rostro-caudal enrichment and decreases antero-ventral markers

WNT signaling has been determined to play multiple developmental roles related to self-renewal and synaptic development, but given its role as a driver of dorsal and caudal patterning in the developing forebrain, we sought to examine how alcohol exposure affected transcription factor expression along the rostral-caudal (A-P) and dorsal-ventral (D-V) neuraxes (O’Leary 2007, Harrison-Uy 2012, Bocchi 2017). It should be noted that among the non-WNT related morphogens be altered with alcohol exposure, nearly all upregulated factors (*FGF2/3, BMP7* and *GLI3*) show specific patterns of expression restricted to the more dorsal and especially caudal aspects of the nascent forebrain (Alzu’bi 2017). As secreted morphogens do not act alone, but in part also coordinate the expression and activity of TFs encoding regional and cell-type specific identities, other factors involved in areal coding were examined in these data (Fig. 6A).

**Figure 6.**
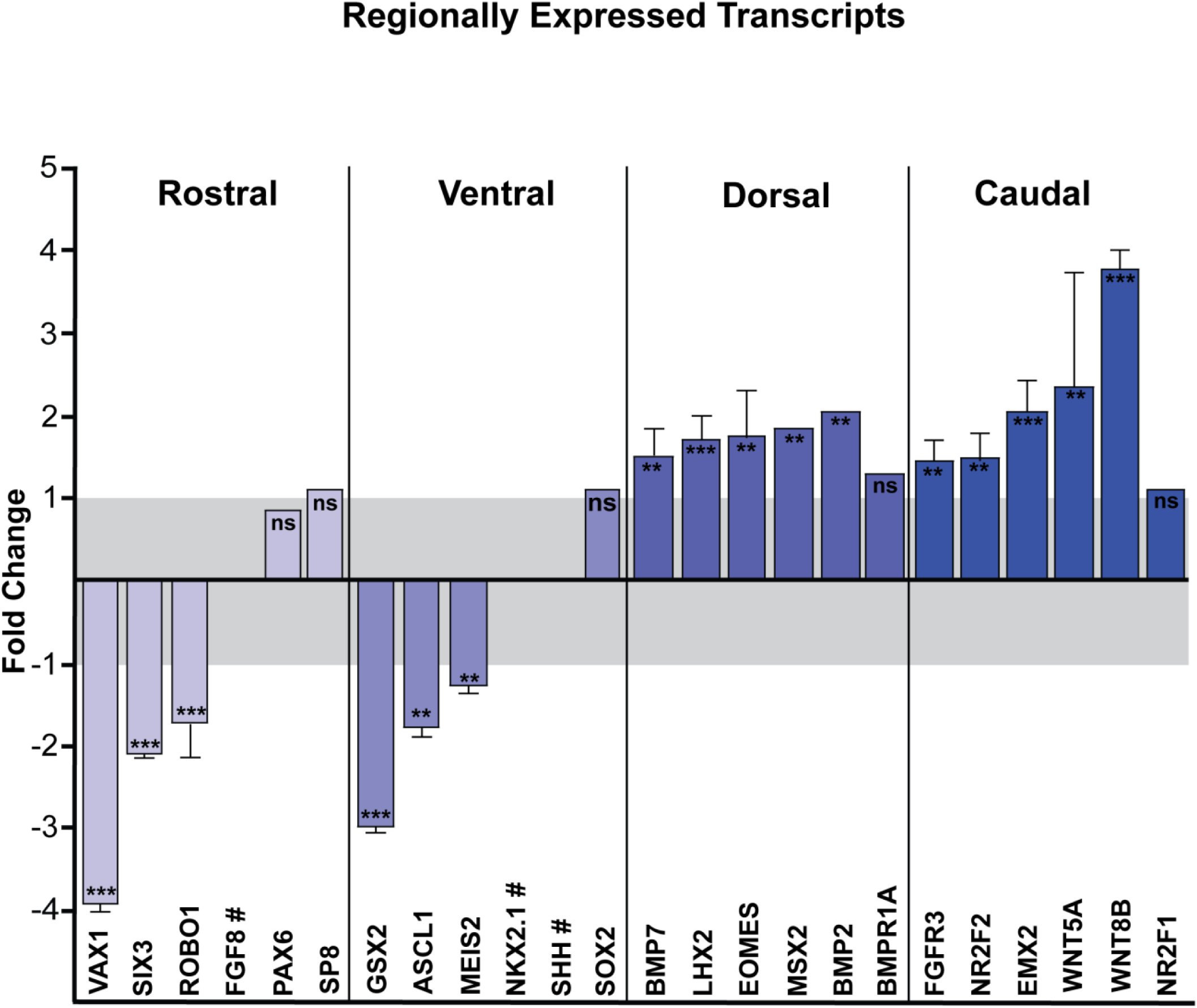

**Figure 7.**
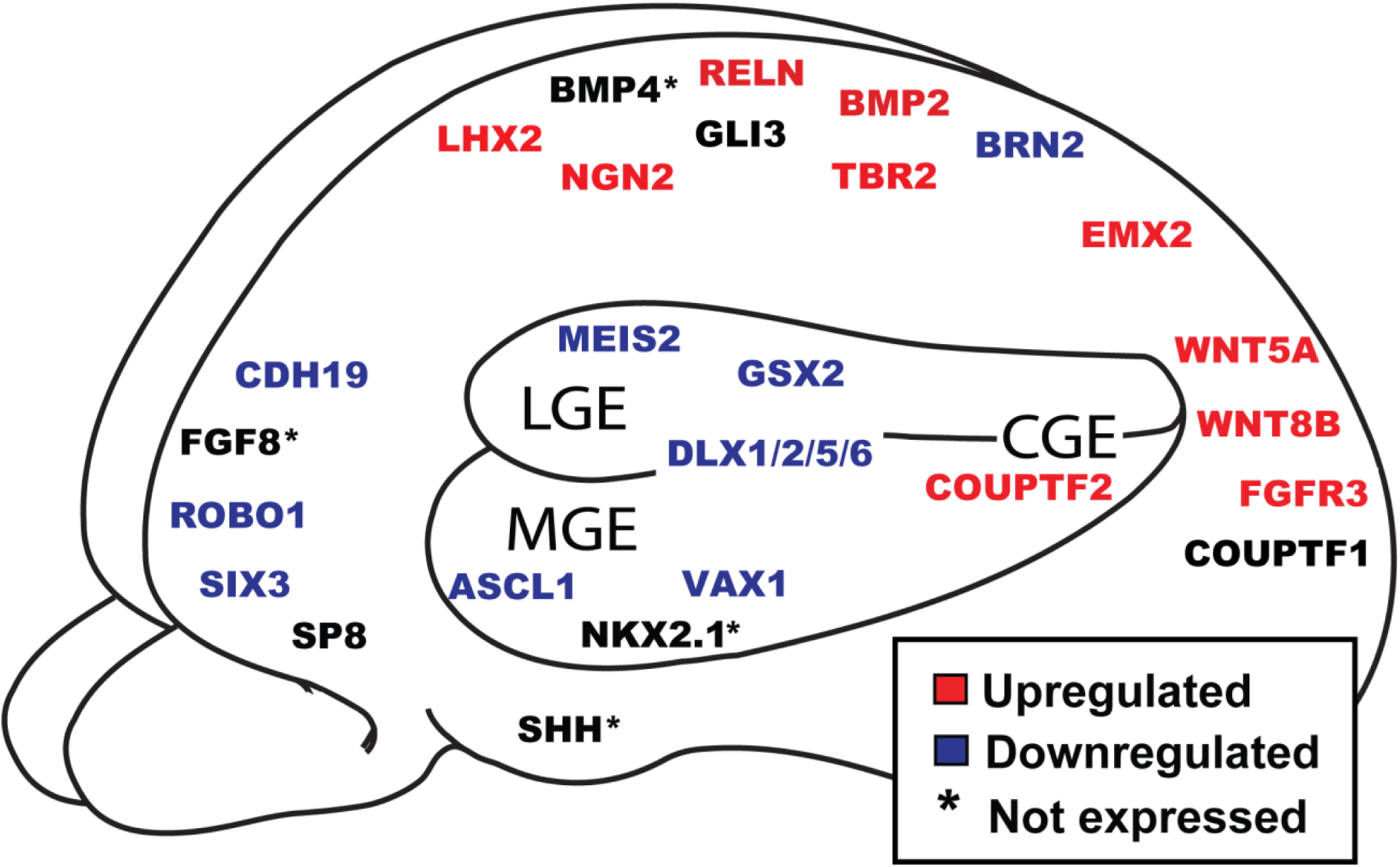

Several important markers of the rostral aspect of the developing forebrain were either not expressed above background levels or were strongly downregulated in context of alcohol exposure. Among these, *FGF8*, a protein understood to coordinate the assembly of the rostral neural ridge (ANR), a crucial signaling center that regulates many regional and cell fate decisions was not found to be expressed in these data. This could indicate that the model described in this report tends to generate neurons of an overall more caudal cortical phenotype, and may complicate drawing conclusions about the regionalizing effects of alcohol. However the downregulation of transcription factors such *ROBO1* and *SIX3*, also known to help establish rostral regional identity, still strongly suggest an alcohol-mediated shift away from whatever rostral character exists within these cells. Additionally, transcription factor ventral-rostral homeobox 1 (*VAX1*), which functions to specify ventral domains of the developing forebrain as well as coordinating midline craniofacial morphogenesis, shows a nearly 4-fold downregulation (Hallonet 1999).

Downregulation of transcripts with ventral patterns of expression was also noted in these data, although perhaps to a lesser degree than purely rostral factors. This effect may be in part due to the significant decrease in transcripts implicated in GABAergic IN development in the ganglionic eminences (GE). Multiple of these factors were significantly decreased with alcohol, including the homeobox genes in the *DLX* family and *GSX2*, as well as *ASCL1* (Fig 6B). The more caudal GE marker *COUPTFII* was however upregulated. As previously mentioned, hedgehog signaling through the smoothened receptor is also understood to be crucial regulator of ventral forebrain specification, but *SHH* expression interesting proved to be well below our threshold of detection. Changes to dorsally-specified genes was more minimal. The transcription factor *MSX2*, which was shown to be upregulated, shares a role with *VAX1* in coordinating craniofacial development, altered development of which is a canonical feature of FAS (Depew 2002, Jeong 2008, Jin 2011, Geohegan 2018).

The majority of the alterations to regionally-restricted genes display a coordinated upregulation of caudally restricted gene products. The changes to caudally expressed *WNTs* drove much of this observation, but TFs such as *EMX2* and *COUPTFII* were also more highly expressed with alcohol exposure. *FGFR3* expression, also increased with alcohol, has been demonstrated to regulate the development of caudal telencephalon and proper migration and integration of GABAergic INs into the cortex (Moldrich 2011). Taken together these data suggest a diminished rostral character of NPCs and neurons exposed to alcohol, with a concomitant upregulation of transcripts associated with caudal forebrain regions suggesting an overall caudalization of the alcohol-exposed forebrain in very early stages of development.

## Discussion

The transcriptomic data reported here provide unique insights into the potential mechanisms of cortical dysfunction in patients with FASD. We found that many of the alcohol-induced changes could be clustered into a relatively small number of biologically relevant categories pertaining to cell fate decisions, synaptic specificity and cortical regionalization. Consistent with previous reports, WNT signaling, known to be critical in multiple aspects of development such as self-renewal and cell fate commitment, proved to be disproportionately affected by the presence of alcohol compared to other secreted morphogens and mitogens. Based on the role of WNT signaling in determination of areal identity, we examined the regulation of a number of genes with regionally restricted patterns of expression and found evidence for a spectrum of up- and down-regulation along the nascent neuraxis. RNA species that are restricted in expression to rostro-ventral aspects of the forebrain in normal development showed notable downregulation, while more dorso-caudal gene products showed concurrent upregulation. Taken together, these data point to a potential primary effect of alcohol on WNT signaling, with the downstream consequence being overall caudalization of developing cerebral cortex. This is consistent with many well characterized teratogenic effects of developmental alcohol exposure, and may provide insights into a previously underappreciated genetic FASD phenotype with important implication for the functional deficits observed in these clinical populations.

Interesting among these data was the increase in overall gene expression, with more than twice as many genes upregulated in the context of alcohol exposure versus control cultures. This finding is consistent however with recent work concerning the epigenetic effects of blood-brain barrier permeable acetyl groups, generated by the metabolism of alcohol. A report from the University of Pennsylvania demonstrates the direct acetylation of histones in the gestating mouse brain from the maternal consumption of isotope-labeled alcohol (Mews 2019). This alcohol-mediated brain histone acetylation in the embryo would lead to overall relaxing of the chromatin structure and an increase in gene expression, in agreement with these findings.

Although we were surprised to observe in this work specific alterations in transcripts associated with glutamatergic specification and synaptogenesis this has been previously observed by other groups. Phenotypic evidence for this imbalances from clinical studies suggests that individuals with FASD display significantly increased rates of seizure disorders compared to non-exposed individuals (Nicita 2014). Rodent models exposed to comparable alcohol concentrations to those used in this report demonstrate increased rates of glutamatergic differentiation and specification, which is hypothesized to proceeded in a *PAX6*-dependent manner (Kim 2010). Interestingly, despite high expression of *PAX6* in the vast majority of neural precursors generated in this differentiation scheme, we found no difference in *PAX6* expression with alcohol exposure in the current study, nor in our previous report, suggesting an alternative mechanism of E/I compensation (Larsen 2016). While multiple TFs thought to be required for nearly all GABA IN fates were downregulated (e.g. *ASCL1, GSX2, DLX1/2/5/6*), studies with DLX1/2 knockout mice, despite leading to an embryonic lethal phenotype, surprisingly still demonstrate the generation of GABAergic neurons (Le 2007). Specifically, one possible explanation for this disparity is an upregulation of GABAergic neurons that are less affected by the loss of many upstream regulators thought to be required for GABA IN specification. Single-cell RNA sequencing experiments designed to assess cellular diversity in adult mouse visual cortex identified 49 distinct cell types, 23 of which represented diverse interneuron population, including two novel *NDNF*^+^ IN subtypes, upregulated 2.61-fold with alcohol in these data (Tasic 2016). Perhaps the lack of network-level alterations to frequency and amplitude of miniature psot-synaptic potentials is compensated for not by overall changes to the number of INs, but rather a protective redistribution of their respective molecular identities among the diversity of IN subtypes that the field is only beginning to understand. Another possibility is more subtle alterations to noncoding RNA species such as microRNAs (miRNA), which represent an underappreciated set of potential targets for understanding mechanisms of disease pathology, due to their diverse roles in coordinating convergent signaling pathways (Hébert 2008).

Key among the findings in this report was the preferential dysregulation of WNT signaling with alcohol, an alteration with many potential implications. The secreted family of WNT signaling molecules and their downstream effectors are known to regulate diverse developmental processes from the delineation of the initial germ layers to maintenance of stem cell proliferation and overall patterning of the forebrain (Lindsley 2006, Harrison-Uy 2012, Merrill 2012). Additionally, WNTs and their receptors have been found to influence synaptogenesis and synapse strengthening/maintenance (Ahmad-Annuar 2006). WNT ligands necessary for cell cycle regulation (WNT1/3A) were not affected by alcohol exposure, but additional family members with influence on synaptogenesis did show surprising degrees of dysregulation in these data. For instance, the highly expressed WNT7A ligand promotes synaptic assembly from the presynaptic bouton as well as synaptic vesicle recycling (Hall 2000). Although WNT7A ligand expression was unaltered with alcohol, shRNA knockdown of the receptor protein frizzled5 (FZD5) suggests that FZD5 is necessary for WNT7A’s activity at nascent synapses (Sahores 2010). Interestingly, FZD5 is one of the only WNT-related transcripts to show significant reduction with alcohol exposure. This suggests that despite WNT7A not being regulated directly by alcohol, its effect through the FZD5 receptor is likely compromised, potentially reducing WNT7A’s positive effect on excitatory glutamatergic tone overall as one potential mechanism of E/I compensation through WNT signaling. Another WNT protein that has been extensively studied for its role in synapse formation and maintenance is WNT5A – among the most highly upregulated in our study (Salinas 2012). In a FZD5-independent manner, WNT5A promotes synaptic assembly through binding Ror tyrosine receptors, which are also highly upregulated (Paganoni 2010). Interestingly, WNT5A increases the clustering of PSD95 and GABAA receptors when applied to hippocampal neurons in vitro, increasing mini inhibitory postsynaptic current (mIPSC) amplitude but not frequency (Farias 2009, Cuitino 2010). While this does not directly explain the E/I compensation observed, it does clearly indicate the convergence of multiple WNT proteins with alcohol-regulated expression on GABA- and glutamatergic tone rather than cell cycle and proliferation.

In addition to WNT proteins’ roles in synapse formation and maintenance, the preponderance of literature on WNT signaling focuses on their involvement with nervous system patterning (reviewed in Mulligan 2012). Specifically, one WNT family member, WNT5A, has been demonstrated in knockout models to be necessary for the development of caudal brain structures (Kumawat 2016). Broadly speaking however, WNTs tend to antagonize the ventralizing actions of sonic hedgehog in the neural tube, as well as FGF8 signaling in the rostral pole of the telencephalon (Danesin 2009). Expression of FGF8 at the rostral pole of the developing telencephalon is sufficient to coordinate the assembly of an entire regional signaling center referred to as the rostral neural ridge (ANR), (Shanmugalingam 2000). Interestingly, neither SHH nor FGF8 were expressed to detectable levels in neuroepithelial cells derived from the H9 cell line when differentiated via default methods (Supplemental Table 1, Floruta et al., 2017). The lack of FGF8 and SHH expression in these data is interesting as it potentially indicates an inherently caudal cell type due to either the H9 cell line or the default differentiation protocol that is enhanced by alcohol exposure. It is possible that other e/iPSC cell lines that express these factors to a greater basal extent could buffer the phenotype more effectively against caudalizing WNT signaling pathways.

Beyond to WNT signaling, TFs such as NR2F2 (COUP-TFII) and EMX2 are known to coordinate regional cortical identity, as knockout of these factors demonstrably expands the extent of the caudal forebrain rostrally (Mallamaci 2000). These current data were obtained in a human developmental model system however, so it is important to review the emerging work concerning regional distribution of TFs and morphogens in *in utero* human cortex. The Clowry lab in Newcastle, UK has demonstrated that in human developing cortex, the genes for NR2F2 and FGFR3 are among the most significantly enriched in caudal forebrain and both show significant upregulation with alcohol (Alzu’bi 2017). The same study showed that the DLX family, along with ASCl1 and ROBO1 to be among the most rostrally enriched. All of these were downregulated by alcohol exposure in this study. ROBO1, downregulated here with alcohol, has been shown to suppress WNT signaling in human NPCs, potentially further exacerbating the alcohol-mediated caudalization (Andrews 2006). Furthermore, SP8 transcriptional repression by EMX2 has been reported to underlie some of the earliest A-P patterning in developing mouse brain (Sahara 2007) and knockout of SP8 expands caudal CoupTFI/II expression into more rostral cortical regions (Zembryzcki 2007). SP8 was unaltered in these data, but the upregulation of EMX2 would suggest an expansion of cells expressing that marker at the expense of SP8^+^ rostral cell types. Taken together, these data suggest that TFs, in addition to morphogens like WNTs, are coordinating a caudalized cortical phenotype with alcohol exposure.

One final potential caveat we must consider in the discussion of these data is the growing understanding that despite universal expression of certain markers of stemness, various human embryonic stem cell lines have diverse genotypes and epigenetic modifications that likely subtly prime cells of different lines down slightly different developmental trajectories. Genetic diversity among hES cell lines have been appreciated since the early 2000s, and more recently these differences have been investigated more thoroughly by The International Stem Cell Initiative, but the implication for line-specific variability in default patterned cells remains poorly understood (Allegrucci 2006, The International Stem Cell Initiative 2007). More recent investigations into these differences have linked the genetic variation in part to modifications between the chromatic architecture of various pluripotent cell lines, arguing that this lineage-specific variability may lead to differences in terminal differentiation programs at the transcriptomic level (Rubin 2017). This becomes a crucial point when discussing conclusions about the effects of alcohol on cortical regionalization, as these programs may vary between pluripotent lineages and in order to fully verify this caudalizing effect of developmental alcohol exposure on forebrain, multiple ES cell lines must be validated.

## Materials and Methods

### hPSC Maintenance and Differentiation

Neurons were differentiated from the WA09 (H9) pluripotent stem cell line maintained in mTesr1 medium (Stem Cell Technologies, Vancouver, BC, Canada). Stem cells were plated in 6-well plates coated in Matrigel in feeder-free conditions with daily media changes and split 1:6 one day before reaching confluency to ensure maintenance of pluripotency. On day 0 of neuronal differentiation, plated H9 cells were treated with 1mg/ml dispase solution for 5 minutes and lifted into 3D culture as Serum Free Embryoid Bodies (SFEB). SFEBs were maintained for 3 days in mTesr1 media, then transitioned to NSM, continuing with daily media changes. Following 21 days of culture, neurospheres are plated onto glass coverslips treated with poly-D-ornithine (0.1mg/ml, Sigma) and laminin (5ug/ml, Life Tech) in a 24-well format and allowed to adhere. Following adherence of the neurospheres to the coverslips, cells were transitioned to neural differentiation media (NDM) which they were fed every other day and consisted of DMEM/F12 media (ThermoFisher Scientific), supplemented with BDNF and GDNF (10ng/ml; Peprotech, Rocky Hill, NJ), cAMP (1μM; Sigma), ascorbic acid (200μM; Sigma) and laminin (500ng/ml). For alcohol treatment, alcohol was supplemented into cell culture media daily up to 50mM from day 0 until cells were processed for RNA at day 50.

### RNA Isolation and cDNA Prep

Following harvest of day 50 neurons in ice cold PBS solution, RNA species were purified with the miRNeasy RNA Isolation Kit (Qiagen) according to the manufacturer’s recommendations. RNA concentration and quality were assessed with the Nanodrop 2000 spectrophotometer (Thermofisher Scientific). In order to analyze mRNA species in our sample, 100-400ng of total RNA was processed using the SuperScript IV First-Strand Synthesis System (Thermofisher Scientific). miRNA processing utilized the Taqman miRNA Reverse Transcription Kit to synthesize cDNA for analysis (Thermofisher Scientific).

### RNA-sequencing

RNA libraries were prepared and sequenced as described previously (Brown 2017). Briefly, Total RNA was ribo-depleted with Low Input RiboMinus Eukaryote System v2 (Thermo Fisher Scientific, A15027). Ion Total RNA-Seq Kit v2 was used to make cDNA, add barcodes, and amplify the library. RNA libraries were sequenced on P1v2 chips using the Ion Proton(tm) System (Thermo Fisher Scientific, #4476610). Sequencing was completed by the Analytical and Translational Genomics Shared Resource at the University Of New Mexico Cancer Center. Exon feature counts were generated with HTSeq-count (Anders 2014) using a modified RefSeq references, and gene level expression counts were generated by summing exon based counts.

### Nanostring miRNA profiling

Mature miRNA profiling was performed with the NanoString nCounter miRNA Expression Assay Kit at the University of Arizona Genetics Core as previously described. nSolver NanoString software was utilized to calculate the geometric means of all miRNAs and normalize the dataset for analysis. The top 400 most highly expressed miRNAs were selected for further analysis.

### Data Analysis

Raw RNA-Seq reads were aligned to the human genome (CRGh37; hg19) using Torrent Mapping Alignment Program (TMAP, v4.06) and the read counts at gene level were summarized using HT-Seq as previously described [K1]. Only the protein coding genes with at least two samples from one of the conditions whose counts were greater than 10 were retained for the analysis. The read counts data were normalized using trimmed mean of M-values (TMM) implemented in software edgeR[K2][K3]. The genes that are differentially expressed between alcohol and Control were identified using Quasi-likelihood F-test under the generalized linear model framework (also implemented in edgeR), which appropriately took into accounts the correlation between the paired samples. Adjustments for multiple comparisons were conducted via Benjamini-Hochberg false discovery rate (FDR) method [K4] and the significance level was set at FDR = 0.05. Follow-up pathway analyses were performed using DAVID bioinformatics suit (v6.8), software IPA[K5] and Bioconductor package Signaling Pathway Impact Analysis (SPIA)[K6].

Categorizations by gene ontology were carried out using the DAVID Bioinformatics Resources (v6.8) hosted by the Laboratory of Human Retrovirology and Immunoinformatics (LHRI). Pathway analyses performed using the Kyoto Encyclopedia of Genes and Genomes (KEGG) database as well as using Ingenuity Pathway Analysis software (IPA; Qiagen, Hilden, Germany). RNA-sequencing data files have been deposited in the NCBI Sequence Read Archive (SRA) repository (http://www.ncbi.nlm.nih.gov/sra/) with accession number XYZ. Error bars relating to fold change differences represents SEM of calculated fold-change between three biological replicates.

This research was partially supported by UNM Comprehensive Cancer Center Support Grant NCI P30CA118100 and made use of the Analytical and Translational Genomics Shared Resource, which receives additional support from the State of New Mexico.

