## Supplemental Table 5 for "Transcriptomic changes due to early, chronic alcohol exposure during cortical development implicate regionalization, cell-type specification, synaptogenesis and WNT signaling as primary determinants of fetal alcohol Spectrum Disorders"

Analysis: 181212 IPA list FC - mammals, observed only Transcription

Printed: December 12, 2018

IPA Build version: 484108M

| Upstream Regul... | Expr Fold Change | Molecule Type | Predicted Activatio... | Activation z-score | Flags | p-value of ove... | Target molecule... | Mechanistic Netwo... |
|---|---|---|---|---|---|---|---|---|
| POU4F2 | | transcription regulator | | -0.113 | | 4.22E-20 | ↑BARHL2, ↑BC... ...all 15 | |
| ASCL1 | ↓-1.732 | transcription regulator | | 0.901 | bias | 1.10E-17 | ↓CACNA2D3, ↓... ...all 18 | |
| PTF1A | | transcription regulator | | 0.286 | bias | 1.25E-13 | ↓CACNA2D3, ↑... ...all 13 | |
| EOMES | ↑1.738 | transcription regulator | | -0.859 | | 1.37E-12 | ↑BHLHE22, ↑C... ...all 15 | |
| SOX2 | | transcription regulator | | -0.116 | | 2.36E-11 | ↓ASCL1, ↑CXC... ...all 22 | 28 (3) |
| GSX2 | ↓-2.892 | transcription regulator | Inhibited | -2.646 | | 3.24E-11 | ↓ASCL1, ↓DLX1, ... ...all 7 | |
| NEUROG3 | | transcription regulator | | -0.986 | bias | 4.05E-11 | ↓DLX6, ↓EBF1, ... ...all 12 | |
| CTNNB1 | | transcription regulator | | 1.628 | bias | 1.59E-10 | ↑BCL2, ↑BMF, ... ...all 32 | 67 (7) |
| SHH | | peptidase | | 0.376 | bias | 1.50E-09 | ↓ASCL1, ↑BCL2, ...all 15 | |
| bexarotene | | chemical drug | | 1.660 | | 4.95E-09 | ↓ABCA1, ↑BAR... ...all 15 | 15 (2) |
| JAK1/2 | | group | Activated | 3.162 | bias | 6.42E-09 | ↑BHLHE22, ↑K... ...all 10 | |
| SNCA | | enzyme | | 1.324 | | 6.99E-09 | ↑BHLHE22, ↑D... ...all 16 | |
| CREB1 | | transcription regulator | | -1.945 | | 2.73E-08 | ↓ABCA1, ↑BCL2, ...all 24 | |
| POU4F1 | | transcription regulator | | -1.260 | | 4.36E-08 | ↑BCL2, ↑CRAB... ...all 11 | |
| DLX2 | ↓-2.262 | transcription regulator | | -1.038 | | 8.64E-08 | ↓DLX2, ↓DLX5, ... ...all 6 | 6 (2) |
| ZFHX3 | ↓-1.663 | transcription regulator | | | | 7.54E-07 | ↓EBF1, ↑LHX1, ↑... ...all 9 | |
| FOS | | transcription regulator | | -1.897 | | 1.00E-06 | ↓ASL, ↑CASZ1, ... ...all 21 | |
| NOTCH1 | | transcription regulator | | 1.191 | | 1.12E-06 | ↓ASCL1, ↑BCL2, ...all 14 | 15 (3) |
| POU5F1 | | transcription regulator | | 1.630 | | 1.28E-06 | ↓ASCL1, ↑BCL2, ...all 16 | 28 (3) |
| PRAG1 | | kinase | | | | 1.38E-06 | ↑HES1, ↑HEY1, ↑... ...all 3 | |
| HTT | | transcription regulator | | 1.078 | | 1.43E-06 | ↓ABCA1, ↑AD... ...all 24 | 68 (6) |
| thiocoraline | | chemical reagent | Activated | 2.000 | | 1.84E-06 | ↓ASCL1, ↑HES1, ... ...all 4 | |
| tretinoin | | chemical - endogenous... | | 0.738 | | 1.95E-06 | ↓ABCA1, ↓ASC... ...all 39 | 99 (12) |
| BMP4 | | growth factor | | 0.333 | bias | 2.92E-06 | ↓ASCL1, ↑BCL2, ...all 12 | 23 (2) |
| HES1 | ↑1.763 | transcription regulator | | 0.923 | bias | 4.66E-06 | ↓ASCL1, ↑BCL2, ... ...all 7 | |
| PAX6 | | transcription regulator | | -0.071 | | 4.82E-06 | ↓ASCL1, ↑EOM... ...all 12 | 22 (3) |
| RBPJ | | transcription regulator | | 1.516 | bias | 5.24E-06 | ↓ASCL1, ↑BMP7, ...all 10 | 15 (3) |
| TBR1 | ↑2.031 | transcription regulator | | | | 5.47E-06 | ↑FEZF2, ↑NR4A2, ...all 3 | |
| ISL1 | ↓-3.384 | transcription regulator | | -1.414 | | 5.92E-06 | ↑CRABP1, ↑DM... ...all 8 | |
| KLF4 | | transcription regulator | | 1.831 | bias | 6.68E-06 | ↓ASCL1, ↑EMX2, ...all 13 | |
| GDF2 | | growth factor | | 0.000 | bias | 7.02E-06 | ↑COL9A1, ↑CRA... ...all 8 | |
| WNT3A | | cytokine | Activated | 3.378 | bias | 7.58E-06 | ↑BCL2, ↑CEMIP2 ...all 13 | 58 (6) |
| TGFB1 | | growth factor | Activated | 2.438 | | 7.59E-06 | ↓ABCA1, ↑AD... ...all 41 | 99 (11) |
| APP | | other | | -0.500 | | 7.89E-06 | ↓ABCA1, ↓ASC... ...all 26 | 76 (11) |
| GLI3 | | transcription regulator | Activated | 2.000 | | 1.08E-05 | ↑BCL2, ↓DLX2, ... ...all 7 | |
| FOXA2 | | transcription regulator | | 0.728 | | 1.18E-05 | ↑CXCR4, ↓ISL1, ... ...all 11 | 13 (2) |
| FEZF2 | ↑2.178 | transcription regulator | | | | 1.36E-05 | ↑BHLHE22, ↑SL... ...all 3 | |
| Histone h3 | | group | | | | 1.81E-05 | ↑ACKR3, ↑BCL2, ...all 17 | |
| DGCR8 | | enzyme | | -0.970 | | 2.17E-05 | ↑CRABP1, ↑GJA1, ...all 6 | 12 (2) |
| NOG | | growth factor | | 0.243 | bias | 2.17E-05 | ↑BMP7, ↓DLX5, ... ...all 6 | 25 (3) |
| ESR2 | | ligand-dependent nucl... | | -0.095 | | 2.95E-05 | ↓ABCA1, ↑BCL2, ...all 16 | 65 (6) |
| EWSR1 | | other | | 0.447 | bias | 3.08E-05 | ↑ACKR3, ↑BCL2, ... ...all 5 | |
| MIB1 | | enzyme | | | | 4.67E-05 | ↑HES1, ↑HEY1, ↑... ...all 3 | |
| GATA4 | | transcription regulator | | 0.666 | | 4.84E-05 | ↓ASCL1, ↑BCL2, ...all 10 | |
| FEV | | transcription regulator | | | | 5.07E-05 | ↑CXCR4, ↑NFIA, ... ...all 7 | |
| CEBPA | | transcription regulator | | -0.480 | | 5.08E-05 | ↓ASCL1, ↓ASL, ... ...all 16 | 51 (3) |
| BDNF | | growth factor | | -0.006 | | 5.17E-05 | ↑ARPP21, ↑BCL2 ...all 13 | 52 (6) |
| CXCR4 | ↑1.627 | G-protein coupled rece... | | 1.224 | bias | 5.54E-05 | ↑ACKR3, ↑BCL2, ... ...all 6 | |
| DUOXA1 | | other | | | | 7.41E-05 | ↑NEUROD1, ↑N... ...all 3 | |
| FGF8 | | growth factor | | -0.100 | | 7.81E-05 | ↑COL18A1, ↓DL... ...all 7 | |
| PITX2 | | transcription regulator | | 0.310 | | 7.83E-05 | ↑DACH1, ↓DLX2, ...all 8 | 57 (6) |
| BCL11B | | transcription regulator | | | | 7.99E-05 | ↑BCL2, ↑LEF1, ↑... ...all 6 | |
| ACVR1 | | kinase | | | | 8.00E-05 | ↓DLX5, ↑EDIL3, ... ...all 5 | |
| SOX7 | | transcription regulator | | -1.342 | bias | 9.03E-05 | ↓ASCL1, ↑EMX2, ... ...all 5 | |
| NANOG | | transcription regulator | | | | 1.03E-04 | ↓DLX1, ↓DLX5, ↑... ...all 8 | |
| WNT1 | | cytokine | | 1.026 | bias | 1.09E-04 | ↑GAS1, ↑HES1, ↑... ...all 8 | 20 (3) |
| FOXQ1 | | transcription regulator | | | | 1.24E-04 | ↑DACH1, ↓NRX... ...all 2 | |
| HES7 | | transcription regulator | | | | 1.24E-04 | ↑HES1, ↑HEY2 ...all 2 | |
| gamma-secretase inhibit | | chemical reagent | | | | 1.24E-04 | ↑HES1, ↑HEY1 ...all 2 | |
| NOTCH2 | | transcription regulator | | 1.195 | | 1.27E-04 | ↓ASCL1, ↑BCL2, ... ...all 5 | |
| ATN1 | | transcription regulator | | | | 1.35E-04 | ↓EBF1, ↓GAD1, ↑... ...all 8 | |
| miR-141-3p (and other m | | mature microRNA | | -0.587 | bias | 1.43E-04 | ↑DACH1, ↓DLX5, ...all 6 | |
| PAX1 | | transcription regulator | | -1.342 | bias | 1.58E-04 | ↓ASCL1, ↑EMX2, ... ...all 5 | |
| SOX1 | | transcription regulator | Inhibited | -2.236 | bias | 1.67E-04 | ↑EOMES, ↑HES1, ... ...all 6 | |
| LHX1 | ↑2.779 | transcription regulator | | -0.378 | bias | 1.69E-04 | ↓EHD2, ↓FZD5, ... ...all 7 | |
| GRB2 | | kinase | | -1.342 | bias | 1.75E-04 | ↓ASCL1, ↑EMX2, ... ...all 5 | 45 (3) |
| LHX2 | ↑1.700 | transcription regulator | | | | 1.88E-04 | ↑FOXJ1, ↑NEUR... ...all 4 | |
| Tcf7 | | transcription regulator | | | | 1.88E-04 | ↑BCL2, ↑EOMES, ... ...all 4 | 64 (9) |
| ID3 | | transcription regulator | Inhibited | -2.412 | | 1.92E-04 | ↑BCL2, ↑BMP7, ... ...all 9 | |
| FOXD1 | | transcription regulator | | | | 2.13E-04 | ↑FOXJ1, ↓ISL1, ↑... ...all 3 | |
| LNX2 | | other | | | | 2.13E-04 | ↑BCL2, ↑HES1, ↑... ...all 3 | |
| baicalin | | chemical - endogenous... | | | | 2.14E-04 | ↓ASCL1, ↑BCL2, ... ...all 5 | 55 (6) |
| ATOH1 | | transcription regulator | | | | 2.18E-04 | ↑BARHL2, ↓CRH, ...all 4 | |
| SOX3 | | transcription regulator | Inhibited | -2.449 | bias | 2.40E-04 | ↑EOMES, ↑TNF... ...all 6 | |
| GATA3 | | transcription regulator | | 0.333 | | 2.50E-04 | ↓ASCL1, ↑BCL2, ...all 10 | 51 (4) |
| lenalidomide | | chemical drug | | 1.455 | | 2.55E-04 | ↓FZD5, ↑LEF1, ↑... ...all 9 | |
| PRDM8 | ↑2.711 | transcription regulator | | | | 2.82E-04 | ↑BHLHE22, ↑EBF3 ...all 3 | |
| tetrachlorodibenzodioxir | | chemical toxicant | | 1.352 | | 3.20E-04 | ↑BCL2, ↑COL1... ...all 13 | 27 (4) |
| SOX10 | | transcription regulator | | | | 3.32E-04 | ↑ACKR3, ↑LAMB1 ...all 4 | |



©2000-2018 QIAGEN. All rights reserved.

| Upstream Regul... | Expr Fold Change | Molecule Type | Predicted Activatio... | Activation z-score | Flags | p-value of ove... | Target molecule... | Mechanistic Netwo... |
|---|---|---|---|---|---|---|---|---|
| N-[N-(3,5-difluorophena | | chemical - protease inh... | | -1.188 | bias | 3.32E-04 | ↑HES1, ↑HEY1, ↑... ...all 4 | 33 (7) |
| REST | | transcription regulator | | -0.747 | bias | 3.34E-04 | ↓ASCL1, ↓CRH, ... ...all 8 | |
| GATA6 | | transcription regulator | | -0.376 | | 3.35E-04 | ↓ASCL1, ↑EMX2, ... ...all 9 | |
| GMNN | | transcription regulator | Inhibited | -2.449 | bias | 3.37E-04 | ↑EOMES, ↑FOXJ1, ...all 6 | |
| ADCYAP1 | | other | | 0.000 | | 3.40E-04 | ↑BCL2, ↓CACN... ...all 10 | 34 (4) |
| FOXA1 | | transcription regulator | | -0.128 | | 3.49E-04 | ↑COL18A1, ↓ISL1,...all 8 | |
| 6-hydroxydopamine | | chemical toxicant | | -1.083 | bias | 3.60E-04 | ↑BCL2, ↓CRH, ↓... ...all 6 | 66 (8) |
| pyridaben | | chemical toxicant | | 0.447 | | 3.69E-04 | ↑DMRTA2, ↑KIA... ...all 5 | |
| LHX5 | ↑2.388 | transcription regulator | | | | 3.70E-04 | ↓GAD1, ↓SLC32... ...all 2 | |
| FREM2 | ↑1.648 | other | | | | 3.70E-04 | ↑FREM1, ↑FREM2 ...all 2 | |
| OLIG1 | | transcription regulator | | | | 3.70E-04 | ↓DLX1, ↓DLX2 ...all 2 | |
| Rbpjl2 | | other | | | | 3.70E-04 | ↑HEY1, ↑HEY2 ...all 2 | |
| NGF | | growth factor | | -0.068 | bias | 3.75E-04 | ↓ASCL1, ↑BCL2, ... ...all 9 | 83 (12) |
| estrogen receptor | | group | | 0.711 | | 3.76E-04 | ↓ABCA1, ↑BCL2, ...all 10 | 50 (4) |
| lithium | | chemical drug | | 1.016 | | 3.84E-04 | ↑ADCY2, ↑ADR... ...all 6 | |
| haloperidol | | chemical drug | | 0.000 | bias | 4.08E-04 | ↑BCL2, ↓CRH, ↑... ...all 6 | 26 (2) |
| TREM1 | | transmembrane receptor | | 1.131 | bias | 4.19E-04 | ↑ACKR3, ↑BCL2, ... ...all 9 | 69 (7) |
| DICER1 | | enzyme | | -1.039 | | 4.20E-04 | ↑COL18A1, ↓C... ...all 11 | 17 (2) |
| NPY | ↓-2.812 | other | | -1.952 | | 4.28E-04 | ↓CRH, ↓NPY, ↓... ...all 4 | 43 (7) |
| NRXN1 | | transporter | | | | 4.28E-04 | ↑COL4A6, ↓DLX5,...all 4 | |
| beta-estradiol | | chemical - endogenous... | | 0.543 | | 4.57E-04 | ↓ABCA1, ↑ACK... ...all 38 | 96 (12) |
| Betacatenin/TCF | | complex | | | | 4.58E-04 | ↓DLX5, ↑GJA1, ↑... ...all 3 | |
| pyrimidin-2-one beta-rib | | chemical reagent | | | | 4.58E-04 | ↑BCL2, ↓GAD1, ... ...all 3 | |
| ramipril | | chemical drug | | -1.000 | bias | 4.83E-04 | ↑COL4A5, ↓CRH, ...all 4 | |
| CREBBP | | transcription regulator | | -0.152 | bias | 4.99E-04 | ↓ASL, ↑BCL2, ↑... ...all 13 | 55 (5) |
| POLR2A | | enzyme | | | | 5.12E-04 | ↓ASCL1, ↑BMF, ... ...all 5 | |
| Secretase gamma | | complex | | 1.231 | | 5.42E-04 | ↑HES1, ↑HEY1, ↑... ...all 4 | 31 (6) |
| TCF | | group | | | | 5.53E-04 | ↑BCL2, ↓GAD1, ... ...all 6 | 55 (5) |
| RND3 | | enzyme | | | | 5.68E-04 | ↑BCL2, ↑HES1, ↑... ...all 3 | 15 (4) |
| MAML1 | | transcription regulator | | | | 5.68E-04 | ↑HES1, ↑HEY1, ↑... ...all 3 | 15 (3) |
| L-685,458 | | chemical - protease inh... | | | | 5.68E-04 | ↑HES1, ↑HEY1, ↑... ...all 3 | 16 (5) |
| IFNG | | cytokine | | 0.661 | | 5.94E-04 | ↓ABCA1, ↑AD... ...all 29 | 82 (9) |
| FGFR2 | | kinase | | 1.342 | | 6.00E-04 | ↑BCL2, ↓DLX2, ... ...all 7 | |
| triamcinolone acetonide | | chemical drug | | 0.285 | | 6.39E-04 | ↓ANGPTL7, ↑BC... ...all 9 | |
| SATB2 | | transcription regulator | | | | 6.93E-04 | ↑BHLHE22, ↑NE... ...all 3 | |
| metyrapone | | chemical drug | | | | 6.93E-04 | ↓CRH, ↑CXCR4, ... ...all 3 | 34 (4) |
| maneb | | chemical toxicant | | 0.447 | | 6.94E-04 | ↑DMRTA2, ↑KIA... ...all 5 | |
| SMARCB1 | | transcription regulator | Activated | 2.219 | bias | 7.25E-04 | ↑COL18A1, ↑CX... ...all 8 | |
| ID2 | | transcription regulator | | -1.633 | | 7.25E-04 | ↑BCL2, ↑BMP7, ... ...all 8 | |
| SP6 | | transcription regulator | | | | 7.35E-04 | ↑AMBN, ↑LEF1 ...all 2 | |
| DLL3 | | other | | | | 7.35E-04 | ↑HES1, ↑HEY1 ...all 2 | |
| CANX | | other | | | | 7.35E-04 | ↓ABCA1, ↑BCL2 ...all 2 | |
| DLL4 | | other | | 1.131 | | 7.52E-04 | ↑HES1, ↑HEY1, ↑... ...all 4 | 15 (5) |
| JAG1 | | growth factor | | 1.188 | bias | 7.52E-04 | ↑HES1, ↑HEY1, ↑... ...all 4 | 15 (6) |
| APR-246 | | chemical drug | | -1.029 | bias | 8.34E-04 | ↑BCL2, ↓DLX5, ... ...all 4 | |
| SPDEF | | transcription regulator | | -1.342 | bias | 9.83E-04 | ↑COL4A5, ↑COL... ...all 5 | |
| F3 | | transmembrane receptor | Activated | 2.236 | bias | 9.83E-04 | ↑GJA1, ↑HMGA2, ...all 5 | |
| NUMB | | other | | | | 9.94E-04 | ↑HES1, ↑HEY1, ↑... ...all 3 | 31 (5) |
| zoledronic acid | | chemical drug | | | | 1.01E-03 | ↑BCL2, ↑CXCR4, ... ...all 4 | |
| histone deacetylase | | complex | | | | 1.01E-03 | ↑BCL2, ↓CRH, ↑... ...all 4 | |
| CNR1 | | G-protein coupled rece... | | 1.709 | | 1.11E-03 | ↑BMP7, ↓CRH, ↑... ...all 7 | |
| COMMD3-BMI1 | | transcription regulator | | 1.131 | bias | 1.11E-03 | ↓EBF1, ↑LEF1, ↑... ...all 4 | |
| TLX3 | | transcription regulator | | | | 1.17E-03 | ↓ASCL1, ↑NEUR... ...all 3 | |
| FOXN4 | | transcription regulator | | | | 1.22E-03 | ↑NEUROD1, ↑N... ...all 2 | |
| LRP8 | | transmembrane receptor | | | | 1.22E-03 | ↓ABCA1, ↑RELN ...all 2 | |
| CDK2AP1 | | other | | | | 1.33E-03 | ↓FZD5, ↑WNT5A, ...all 4 | 46 (3) |
| DTNBP1 | | other | | | | 1.37E-03 | ↓GAD2, ↓SLC32... ...all 3 | |
| naloxone | | chemical drug | | | | 1.37E-03 | ↑BCL2, ↓CRH, ↓... ...all 3 | 66 (7) |
| MITF | | transcription regulator | Activated | 2.433 | bias | 1.45E-03 | ↑BCL2, ↑HES1, ... ...all 9 | |
| advanced glycation end- | | chemical - endogenous... | | | | 1.45E-03 | ↓ABCA1, ↑BCL2, ... ...all 4 | |
| 2,4,5,2',4',5'-hexachlorob | | chemical toxicant | | 0.000 | | 1.56E-03 | ↑BCL2, ↓EBF1, ↑... ...all 6 | |
| GLIS3 | | transcription regulator | | | | 1.58E-03 | ↓ISL1, ↑NEUROD1...all 3 | |
| CDK8 | | kinase | | | | 1.58E-03 | ↑CRABP1, ↑CXC... ...all 3 | |
| tetracycline | | chemical drug | | -0.577 | | 1.62E-03 | ↑CACNA2D1, ↑... ...all 5 | |
| progesterone | | chemical - endogenous... | | 0.619 | | 1.64E-03 | ↑ADRA2A, ↑BC... ...all 16 | 96 (11) |
| Rxr | | group | | | | 1.79E-03 | ↓ABCA1, ↑BAR... ...all 6 | |
| lithium chloride | | chemical drug | | 1.969 | bias | 1.79E-03 | ↑BCL2, ↑BMP7, ... ...all 6 | 68 (8) |
| Ap2 alpha | | group | | | | 1.81E-03 | ↑BCL2, ↓PTGDS ...all 2 | |
| RO492909? | | chemical drug | | | | 1.81E-03 | ↑HES1, ↑HEY1 ...all 2 | 16 (5) |
| BCL11A | | transcription regulator | | | | 1.81E-03 | ↑BCL2, ↑TBR1 ...all 2 | |
| THBS2 | | other | | | | 1.81E-03 | ↑HES1, ↑HEY1 ...all 2 | |
| ADCY5 | | enzyme | | | | 1.81E-03 | ↑ADCY2, ↑BCL2 ...all 2 | |
| LHX6 | | transcription regulator | | | | 1.81E-03 | ↑AMBN, ↑LEF1 ...all 2 | |
| sparfosic acid | | chemical drug | | | | 1.81E-03 | ↑BCL2, ↑TP73 ...all 2 | |
| Trp53cor1 | | other | | | | 1.81E-03 | ↑LHX5, ↑NPTX1, ... ...all 3 | |
| CDKN1B | | kinase | | -0.758 | bias | 1.95E-03 | ↑BCL2, ↑LEF1, ↑... ...all 6 | |
| SOX2-OCT4-NANOG | | complex | | | | 2.07E-03 | ↓ISL1, ↑LHX5, ↓... ...all 3 | |
| corticosterone | | chemical - endogenous... | | -0.141 | | 2.13E-03 | ↑BCL2, ↓CRH, ↓... ...all 6 | 45 (6) |
| SMO | | G-protein coupled rece... | | | | 2.15E-03 | ↑BCL2, ↓DLX2, ... ...all 5 | 56 (6) |



©2000-2018 QIAGEN. All rights reserved.

| Upstream Regul... | Expr Fold Change | Molecule Type | Predicted Activatio... | Activation z-score | Flags | p-value of ove... | Target molecule... | Mechanistic Netwo... |
|---|---|---|---|---|---|---|---|---|
| SMO | | G-protein coupled rece... | | | | 2.1[illegible]E-03 | ↑BCL2, ↓DLX2, ↑... ...all 5 | 36 (6) |
| BMP2 | | growth factor | | -0.664 | bias | 2.22E-03 | ↓ASCL1, ↑BCL2, ......all 8 | |
| FFAR3 | | G-protein coupled rece... | | 0.447 | | 2.27E-03 | ↓ADGRV1, ↑PCS... ...all 5 | |
| CNTF | | cytokine | | 1.406 | | 2.27E-03 | ↑GJA1, ↑NGFR, ......all 5 | |
| MECP2 | | transcription regulator | | 1.067 | | 2.32E-03 | ↓CRH, ↓DLX5, ↓... ...all 6 | |
| risperidone | | chemical drug | | | | 2.35E-03 | ↑BCL2, ↑NGFR, ......all 3 | |
| COL4A3 | | other | | | | 2.35E-03 | ↑COL4A5, ↑ITGA2...all 3 | |
| NTRK1 | | kinase | | | | 2.35E-03 | ↑BCL2, ↓SCN9A, ......all 3 | |
| DLX1 | ↓-2.288 | transcription regulator | | | | 2.52E-03 | ↓DLX5, ↓DLX6 ...all 2 | |
| RAX | | transcription regulator | | | | 2.52E-03 | ↑HES1, ↑OTX2 ...all 2 | |
| ITGB1BP1 | | transporter | | | | 2.52E-03 | ↑HEY1, ↑HEY2 ...all 2 | 15 (4) |
| TSPO | ↑1.753 | transmembrane receptor | | | | 2.52E-03 | ↑BCL2, ↑TSPO ...all 2 | |
| EPHB4 | | kinase | | 1.000 | bias | 2.52E-03 | ↑BMP7, ↑CXCR4, ...all 4 | |
| BRD2 | | kinase | | | | 2.52E-03 | ↑BCL2, ↑EBF3, ↑... ...all 4 | |
| ACKR3 | ↑1.980 | G-protein coupled rece... | | | | 2.64E-03 | ↑BCL2, ↑COL9A1, ...all 3 | |
| PCYT1A | | enzyme | | | | 2.64E-03 | ↓ABCA1, ↑CDK18, ...all 3 | |
| K-252 | | chemical - kinase inhibi... | | | | 2.64E-03 | ↑AMBN, ↑SLC1... ...all 3 | |
| Hdac | | group | | 1.195 | | 2.79E-03 | ↑BCL2, ↓CRH, ↑... ...all 7 | 66 (8) |
| CYR61 | | other | Activated | 2.219 | bias | 2.79E-03 | ↑ITGA2, ↑MSX1, ......all 5 | |
| OGA | | enzyme | | 0.302 | | 2.96E-03 | ↑BCL2, ↓CACN... ...all 12 | |
| HDAC4 | | transcription regulator | | | | 2.96E-03 | ↑CACNA2D1, ↑... ...all 6 | |
| acetylcholine | | chemical - endogenous... | | | | 2.96E-03 | ↓CRH, ↓NPY, ↑... ...all 3 | 31 (4) |
| Ca2+ | | chemical - endogenous... | | 1.744 | bias | 3.07E-03 | ↑BCL2, ↑CXCR4, ......all 9 | 85 (12) |
| Salmonella enterica sero | | chemical toxicant | Activated | 2.630 | bias | 3.08E-03 | ↑ACKR3, ↑HES1, ......all 7 | |
| NOTCH3 | | transcription regulator | | 1.949 | bias | 3.11E-03 | ↑BCL2, ↑HES1, ......all 4 | |
| 3-nitropropionic acid | | chemical toxicant | | 1.969 | | 3.11E-03 | ↑BCL2, ↑NGFR, ......all 4 | 68 (7) |
| MYC | | transcription regulator | | 1.439 | | 3.16E-03 | ↓ABCA1, ↓ASC... ...all 23 | 61 (7) |
| FGF2 | | growth factor | | 0.988 | bias | 3.25E-03 | ↓ASCL1, ↑BCL2, ...all 11 | 87 (12) |
| NOTCH4 | | transcription regulator | | | | 3.31E-03 | ↑HES1, ↑HEY1, ......all 3 | |
| MED12 | | transcription regulator | | | | 3.33E-03 | ↓ASCL1, ↑NEUR... ...all 2 | |
| OCT4-NANOG | | complex | | | | 3.33E-03 | ↓GSX2, ↑TCF7L1 ...all 2 | |
| GRIP1 | | transcription regulator | | | | 3.33E-03 | ↑FREM1, ↑FREM2...all 2 | |
| naproxen | | chemical drug | | | | 3.33E-03 | ↑COL18A1, ↑NG... ...all 2 | |
| chlorpyrifos | | chemical toxicant | | | | 3.33E-03 | ↓CRH, ↓NPY ...all 2 | |
| SIX1 | | transcription regulator | | | | 3.67E-03 | ↑DACH1, ↓DLX5, ...all 3 | |
| AR | | ligand-dependent nucl... | | -0.199 | bias | 3.69E-03 | ↓ABCA1, ↑BMF, ...all 13 | |
| MSC | | transcription regulator | | 1.000 | | 3.78E-03 | ↑BMP7, ↑IFI44, ↑... ...all 4 | |
| L-dopa | | chemical - endogenous... | | 1.000 | | 3.92E-03 | ↑ARPP21, ↑BCL2 ...all 16 | |
| KN 93 | | chemical - kinase inhibi... | | | | 4.07E-03 | ↓ABCA1, ↓GAD1, ...all 3 | 24 (2) |
| PDLIM2 | | other | | 0.277 | | 4.10E-03 | ↑BCL2, ↑DMRT3, ...all 5 | |
| SMARCA4 | | transcription regulator | | 0.849 | bias | 4.16E-03 | ↓ABCA1, ↑BCL2, ...all 15 | |
| HRAS | | enzyme | | 1.990 | | 4.22E-03 | ↑ADRA2A, ↓AS... ...all 15 | 57 (6) |
| Mek | | group | Inhibited | -2.197 | | 4.24E-03 | ↓ASCL1, ↑BMF, ......all 8 | |
| neomycin | | chemical drug | | | | 4.25E-03 | ↑HES1, ↑HEY1 ...all 2 | |
| phosphatidylcholine | | chemical - endogenous... | | | | 4.25E-03 | ↓ABCA1, ↑BCL2 ...all 2 | |
| Mir218 | | microRNA | | | | 4.25E-03 | ↑CXCR4, ↑LEF1 ...all 2 | |
| (-)-epicatechin gallate | | chemical - endogenous... | | | | 4.25E-03 | ↓DLX5, ↓WWTR1 ...all 2 | |
| SERPINB2 | | other | | | | 4.25E-03 | ↑BCL2, ↑TP73 ...all 2 | |
| BRMS1 | | enzyme | | | | 4.25E-03 | ↑CXCR4, ↑GJA1 ...all 2 | |
| PD168393 | | chemical - kinase inhibi... | | | | 4.25E-03 | ↑BCL2, ↑RGS16 ...all 2 | |
| prodigiosin | | chemical toxicant | | | | 4.25E-03 | ↑BCL2, ↑TP73 ...all 2 | |
| BRAF | | kinase | | | | 4.29E-03 | ↓ASCL1, ↑BMF, ......all 5 | |
| LEP | | growth factor | Activated | 2.239 | | 4.67E-03 | ↓ASL, ↑BCL2, ↓... ...all 12 | 70 (8) |
| RUNX1 | | transcription regulator | | -0.164 | | 4.75E-03 | ↑HEY2, ↑HMGA2, ...all 7 | |
| 2-deoxyglucose | | chemical drug | | -0.228 | | 4.82E-03 | ↑BCL2, ↑BMF, ↓... ...all 4 | 79 (10) |
| estrogen | | chemical drug | | 1.461 | bias | 4.83E-03 | ↓ABCA1, ↑BCL2, ......all 9 | 77 (8) |
| PRKAR1A | | kinase | | | | 4.92E-03 | ↑BCL2, ↑NR4A2, ......all 3 | 30 (4) |
| POSTN | | other | | | | 4.92E-03 | ↑HES1, ↑HEY1, ↑... ...all 3 | |
| SOX4 | | transcription regulator | | 1.671 | bias | 5.03E-03 | ↑ARPP21, ↑BCL2, ...all 7 | |
| AGT | | growth factor | | 1.395 | bias | 5.09E-03 | ↓ASL, ↑BCL2, ↑... ...all 12 | 113 (15) |
| Pdgf (complex) | | complex | | -0.391 | bias | 5.11E-03 | ↑BCL2, ↑GJA1, ↑... ...all 5 | 50 (10) |
| POMC | | other | | 1.172 | | 5.11E-03 | ↓CRH, ↑CXCR4, ......all 5 | 77 (10) |
| CIP2A | | other | | | | 5.11E-03 | ↑BCL2, ↑NPTX1, ......all 4 | |
| pCPT-cAMP | | chemical - kinase inhibi... | | | | 5.28E-03 | ↓ABCA1, ↓SST ...all 2 | |
| NAP1L1 | | other | | | | 5.28E-03 | ↑CXCR4, ↑TTR ...all 2 | |
| NEIL2 | | enzyme | | | | 5.28E-03 | ↓ASCL1, ↑NEUR... ...all 2 | |
| NQO2 | | enzyme | | | | 5.28E-03 | ↑BCL2, ↑CXCR4 ...all 2 | 66 (7) |
| CARTPT | | other | | | | 5.28E-03 | ↓CRH, ↑TRH ...all 2 | 43 (6) |
| RELN | ↑2.486 | peptidase | | | | 5.28E-03 | ↓ABCA1, ↓GAD1 ...all 2 | |
| FOXJ1 | ↑1.825 | transcription regulator | | | | 5.28E-03 | ↓DLX2, ↑FOXJ1 ...all 2 | |
| allopregnanolone | | chemical - endogenous... | | | | 5.28E-03 | ↓CRH, ↑GABRA4 ...all 2 | |
| NFkB (complex) | | complex | | 1.279 | bias | 5.32E-03 | ↑ADAMTS9, ↑... ...all 15 | 71 (7) |
| hymecromone | | chemical drug | | | | 5.38E-03 | ↑ACKR3, ↑BCL2, ......all 3 | |
| mifepristone | | chemical drug | | -0.156 | | 5.52E-03 | ↑BCL2, ↑BMP7, ......all 9 | 56 (5) |
| IL1B | | cytokine | | 0.495 | bias | 5.59E-03 | ↑BCL2, ↑BMF, ......all 19 | 56 (7) |
| LEF1 | ↑2.203 | transcription regulator | | | | 5.72E-03 | ↑BCL2, ↑CXCR4, ......all 4 | |
| IL7 | | cytokine | | 1.029 | bias | 5.87E-03 | ↑BCL2, ↑CXCR4, ......all 6 | |
| kainic acid | | chemical toxicant | | 0.130 | bias | 5.87E-03 | ↑BCL2, ↑CACN... ...all 6 | 58 (6) |
| TFAP2A | | transcription regulator | | | | 6.03E-03 | ↓ABCA1, ↑BCL2, ......all 5 | |



©2000-2018 QIAGEN. All rights reserved.

| Upstream Regul... | Expr Fold Change | Molecule Type | Predicted Activatio... | Activation z-score | Flags | p-value of ove... | Target molecule... | Mechanistic Netwo... |
|---|---|---|---|---|---|---|---|---|
| BCL2L1 | | other | | 1.969 | | 6.05E-03 | ↑BCL2, ↑MSX1, ↑... ...all 4 | |
| NRAS | | enzyme | | -0.659 | | 6.13E-03 | ↑ADCY2, ↑BCL2, ... ...all 7 | |
| INHBA | | growth factor | | 0.000 | bias | 6.30E-03 | ↑BCL2, ↑CXCR4, ... ...all 7 | 96 (12) |
| decitabine | | chemical drug | Activated | 2.249 | bias | 6.31E-03 | ↑CASZ1, ↑CXC... ...all 13 | |
| TXK | | kinase | | | | 6.40E-03 | ↑EOMES, ↑ZBTB... ...all 2 | |
| LB-205 | | chemical reagent | | | | 6.40E-03 | ↑BCL2, ↑NGFR ...all 2 | |
| VEGFC | | growth factor | | | | 6.40E-03 | ↑BCL2, ↑CXCR4 ...all 2 | |
| NEUROD2 | ↑2.233 | transcription regulator | | | | 6.40E-03 | ↑NEUROD2, ↑T... ...all 2 | |
| pterostilbene | | chemical drug | | | | 6.40E-03 | ↑BCL2, ↑HES1 ...all 2 | 15 (4) |
| VEGFA | | growth factor | Activated | 2.398 | bias | 6.42E-03 | ↑BCL2, ↑CXCR4, ... ...all 9 | 70 (7) |
| paclitaxel | | chemical drug | | | | 6.55E-03 | ↑ACKR3, ↑BCL2, ... ...all 9 | |
| AHR | | ligand-dependent nucl... | | 0.481 | | 6.60E-03 | ↑ADAMTS2, ↑... ...all 10 | 51 (3) |
| doxorubicin | | chemical drug | | -0.676 | | 6.78E-03 | ↑BCL2, ↓GAD1, ... ...all 11 | 78 (9) |
| dopamine | | chemical - endogenous... | | 0.200 | | 6.80E-03 | ↑BCL2, ↓CACN... ...all 5 | |
| EZH2 | | transcription regulator | | | | 6.86E-03 | ↑ACKR3, ↑BCL2, ...all 10 | |
| EGR2 | | transcription regulator | | 0.555 | bias | 7.34E-03 | ↑BCL2, ↑CRABP1, ...all 6 | |
| fatty acid | | chemical - endogenous... | | | | 7.34E-03 | ↓ABCA1, ↑BCL2, ... ...all 5 | 66 (7) |
| FANCC | | other | | | | 7.46E-03 | ↑HES1, ↑IFI44L, ↑... ...all 4 | |
| NQO1 | | enzyme | | | | 7.50E-03 | ↑BCL2, ↑CXCR4, ... ...all 3 | |
| dexamethasone | | chemical drug | | -0.942 | | 7.55E-03 | ↑ACKR3, ↑ADA... ...all 35 | 93 (9) |
| highly active antiretrovira | | chemical drug | | | | 7.63E-03 | ↑BCL2, ↑CXCR4 ...all 2 | |
| STAR | | transporter | | | | 7.63E-03 | ↓ABCA1, ↓CRH ...all 2 | |
| TGFBR3 | | kinase | | | | 7.63E-03 | ↑BCL2, ↑COL4A6 ...all 2 | |
| nordihydroguaiaretic aci | | chemical drug | | | | 7.63E-03 | ↑BCL2, ↓CRH ...all 2 | |
| PRNP | | other | | -0.795 | | 7.84E-03 | ↑BCL2, ↑CHST8, ... ...all 4 | 32 (4) |
| MSX2 | | transcription regulator | | | | 8.10E-03 | ↑LEF1, ↓SIX3, ↑... ...all 3 | |
| FOXP1 | | transcription regulator | | | | 8.10E-03 | ↑GAS1, ↑GJA1, ↑... ...all 3 | |
| ZNF217 | ↑1.666 | transcription regulator | | | | 8.65E-03 | ↑EOMES, ↓GAD1, ...all 4 | |
| cilostazol | | chemical drug | | | | 8.72E-03 | ↓ABCA1, ↑BCL2, ... ...all 3 | 50 (6) |
| SGPL1 | | enzyme | | | | 8.95E-03 | ↓ABCA1, ↑BCL2 ...all 2 | |
| LMX1A | ↑2.639 | transcription regulator | | | | 8.95E-03 | ↑HES1, ↑NEURO... ...all 2 | |
| GFAP | | other | | | | 8.95E-03 | ↑ITGA2, ↑KCNJ2 ...all 2 | |
| phencyclidine | | chemical drug | | | | 8.95E-03 | ↑BCL2, ↓NRG1 ...all 2 | |
| topiramate | | chemical drug | | | | 8.95E-03 | ↓CRH, ↑TRH ...all 2 | |
| VCAN | | other | | 0.928 | | 9.06E-03 | ↑ADAMTS9, ↑IFI... ...all 6 | |
| Creb | | group | | 1.109 | bias | 9.07E-03 | ↑BCL2, ↓CRH, ↑... ...all 9 | |
| (+)-MK-801 | | chemical drug | | -0.152 | | 9.07E-03 | ↓CRH, ↑GJA1, ↑... ...all 4 | |
| paraquat | | chemical toxicant | | 0.447 | bias | 9.17E-03 | ↑DMRTA2, ↑KIA... ...all 5 | |
| cyclic AMP | | chemical - endogenous... | | 0.503 | bias | 9.31E-03 | ↓ABCA1, ↑ADR... ...all 8 | 77 (10) |
| NEUROD1 | ↑1.768 | transcription regulator | | | | 9.37E-03 | ↑NEUROD1, ↑N... ...all 3 | |
| NR3C2 | | ligand-dependent nucl... | | 1.982 | | 9.50E-03 | ↑BCL2, ↓CACN... ...all 5 | |
| methamphetamine | | chemical drug | | -0.152 | | 9.51E-03 | ↑BCL2, ↑BMP7, ↓... ...all 4 | |
| RARA | | ligand-dependent nucl... | | -0.308 | | 9.80E-03 | ↓ABCA1, ↓CRH, ↑... ...all 9 | 68 (7) |
| CEACAM1 | | transporter | | | | 1.00E-02 | ↑COL18A1, ↑CX... ...all 3 | |
| TP73 | ↑2.476 | transcription regulator | | -0.193 | | 1.02E-02 | ↑BCL2, ↑BMP7, ... ...all 10 | |
| cetuximab | | biologic drug | | | | 1.04E-02 | ↑BCL2, ↓NRG1 ...all 2 | |
| POU3F2 | ↓-1.670 | transcription regulator | | | | 1.04E-02 | ↓CRH, ↑NEURO... ...all 2 | |
| mir-196 | | microRNA | | | | 1.04E-02 | ↑BCL2, ↑HMGA2 ...all 2 | |
| NHLH2 | ↑2.652 | transcription regulator | | | | 1.04E-02 | ↓ASCL1, ↑TRH ...all 2 | |
| AICDA | | enzyme | | | | 1.04E-02 | ↓ROBO1, ↓ZFHX3 ...all 2 | |
| HNF1B | | transcription regulator | | | | 1.05E-02 | ↑HMGA2, ↑LA... ...all 5 | |
| ethanol | | chemical - endogenous... | | 0.344 | | 1.06E-02 | ↓ABCA1, ↑ACK... ...all 10 | |
| ERBB2 | | kinase | | -0.803 | bias | 1.11E-02 | ↑BCL2, ↑BMP7, ... ...all 16 | |
| Pde1 | | group | | | | 1.12E-02 | ↓CRH ...all 1 | |
| hydroxamic acid | | chemical - other | | | | 1.12E-02 | ↑CXCR4 ...all 1 | |
| Ca2 ATPase | | group | | | | 1.12E-02 | ↑TP73 ...all 1 | |
| Ryr | | group | | | | 1.12E-02 | ↑KCNJ2 ...all 1 | |
| dichlobenil | | chemical toxicant | | | | 1.12E-02 | ↓GAD1 ...all 1 | |
| sodium thiosulfate | | chemical drug | | | | 1.12E-02 | ↑BCL2 ...all 1 | |
| DMRT3 | ↑2.643 | transcription regulator | | | | 1.12E-02 | ↑NEUROG2 ...all 1 | |
| RGN | | enzyme | | | | 1.12E-02 | ↑BCL2 ...all 1 | |
| DLX6-AS1 | ↓-2.137 | other | | | | 1.12E-02 | ↓DLX6 ...all 1 | |
| colestimide | | chemical drug | | | | 1.12E-02 | ↓ABCA1 ...all 1 | |
| NPAS1 | | transcription regulator | | | | 1.12E-02 | ↑RELN ...all 1 | |
| NLGN2 | | enzyme | | | | 1.12E-02 | ↓GAD2 ...all 1 | |
| SPAST | | enzyme | | | | 1.12E-02 | ↑EMX2 ...all 1 | |
| ZG16B | | other | | | | 1.12E-02 | ↑CXCR4 ...all 1 | |
| NTNG1 | | other | | | | 1.12E-02 | ↑LRRC4C ...all 1 | |
| POU5F2 | | transcription regulator | | | | 1.12E-02 | ↓CRH ...all 1 | |
| METAP2 | | peptidase | | | | 1.12E-02 | ↑BCL2 ...all 1 | |
| FERD3L | | transcription regulator | | | | 1.12E-02 | ↑HES1 ...all 1 | |
| Tas2r103 | | other | | | | 1.12E-02 | ↑TRH ...all 1 | |
| FK-962 | | chemical drug | | | | 1.12E-02 | ↓SST ...all 1 | |
| STX12 | | other | | | | 1.12E-02 | ↓ABCA1 ...all 1 | |
| voxtalisib | | chemical drug | | | | 1.12E-02 | ↑BCL2 ...all 1 | |
| FANCG | | other | | | | 1.12E-02 | ↑HES1 ...all 1 | |
| FANCL | | enzyme | | | | 1.12E-02 | ↑HES1 ...all 1 | |
| ACSL1 | | enzyme | | | | 1.12E-02 | ↓ABCA1 ...all 1 | |
| NAPA | | transporter | | | | 1.12E-02 | ↑BCL2 ...all 1 | |



©2000-2018 QIAGEN. All rights reserved.

| Upstream Regul... | Expr Fold Change | Molecule Type | Predicted Activatio... | Activation z-score | Flags | p-value of ove... | Target molecule... | Mechanistic Netwo... |
|---|---|---|---|---|---|---|---|---|
| OTP | | transcription regulator | | | | 1.12E-02 | ↓CRH ...all 1 | |
| PLOD1 | | enzyme | | | | 1.12E-02 | ↑GJA1 ...all 1 | |
| TRAK2 | | transporter | | | | 1.12E-02 | ↑KCNJ2 ...all 1 | |
| mir-592 | | microRNA | | | | 1.12E-02 | ↑NGFR ...all 1 | |
| mir-329 | | microRNA | | | | 1.12E-02 | ↓GAD2 ...all 1 | |
| mir-500 | | microRNA | | | | 1.12E-02 | ↓GAD1 ...all 1 | |
| miR-33-5p (and other mi | | mature microRNA | | | | 1.12E-02 | ↓ABCA1 ...all 1 | |
| miR-383-5p (miRNAs w/ | | mature microRNA | | | | 1.12E-02 | ↑NEUROD1 ...all 1 | |
| ACAT2 | | enzyme | | | | 1.12E-02 | ↓ABCA1 ...all 1 | |
| DYNLT3 | | other | | | | 1.12E-02 | ↑BCL2 ...all 1 | |
| SNTB1 | | other | | | | 1.12E-02 | ↓ABCA1 ...all 1 | |
| SCD5 | | enzyme | | | | 1.12E-02 | ↑WNT7B ...all 1 | |
| NOTCH2NLA/NOTCH2N | | other | | | | 1.12E-02 | ↑HES1 ...all 1 | |
| AP3M2 | | transporter | | | | 1.12E-02 | ↓SLC32A1 ...all 1 | |
| RIT1 | | enzyme | | | | 1.12E-02 | ↑BCL2 ...all 1 | |
| PHF5A | | transcription regulator | | | | 1.12E-02 | ↑GJA1 ...all 1 | |
| FANCE | | other | | | | 1.12E-02 | ↑HES1 ...all 1 | |
| STARD3 | | transporter | | | | 1.12E-02 | ↓ABCA1 ...all 1 | |
| CD164 | | other | | | | 1.12E-02 | ↑CXCR4 ...all 1 | |
| FZD2 | | G-protein coupled rece... | | | | 1.12E-02 | ↑WNT5A ...all 1 | |
| tetramethylpyrazine | | chemical - endogenous... | | | | 1.12E-02 | ↑CXCR4 ...all 1 | |
| CORT-108297 | | chemical reagent | | | | 1.12E-02 | ↓CRH ...all 1 | |
| TPD52 | | other | | | | 1.12E-02 | ↑BCL2 ...all 1 | |
| LMO3 | | other | | | | 1.12E-02 | ↓ASCL1 ...all 1 | |
| CWP232228 | | chemical reagent | | | | 1.12E-02 | ↑LEF1 ...all 1 | |
| melarsoprol | | chemical drug | | | | 1.12E-02 | ↑BCL2 ...all 1 | |
| pegademase bovine | | biologic drug | | | | 1.12E-02 | ↓PDE5A ...all 1 | |
| oxaprozin | | chemical drug | | | | 1.12E-02 | ↑NGFR ...all 1 | |
| gastric acid | | chemical - endogenous... | | | | 1.12E-02 | ↓SST ...all 1 | |
| 2,3-dichloro-5,8-dihydro | | chemical toxicant | | | | 1.12E-02 | ↑TP73 ...all 1 | |
| glycosphingolipid | | chemical - other | | | | 1.12E-02 | ↑AMBN ...all 1 | |
| pregabalin | | chemical drug | | | | 1.12E-02 | ↑CACNA2D1 ...all 1 | |
| polysialic acid | | chemical drug | | | | 1.12E-02 | ↑NGFR ...all 1 | |
| stanozolol | | chemical drug | | | | 1.12E-02 | ↑NGFR ...all 1 | |
| 6alpha-fluorotestosteror | | chemical toxicant | | | | 1.12E-02 | ↑BCL2 ...all 1 | |
| THRB | | ligand-dependent nucl... | | -0.555 | | 1.13E-02 | ↑CXCR4, ↑GSN,↑... ...all 7 | |
| HOXC6 | | transcription regulator | | | | 1.15E-02 | ↑BCL2, ↑BMP7,↑... ...all 3 | |
| RUNX2 | | transcription regulator | | 0.218 | bias | 1.17E-02 | ↑AMBN, ↑BCL2,... ...all 5 | |
| APBB1 | | transcription regulator | | | | 1.19E-02 | ↑HES1, ↑TP73 ...all 2 | |
| miR-128-3p (and other n | | mature microRNA | | | | 1.19E-02 | ↑AFF1, ↑RELN ...all 2 | |
| SPINT1 | | other | | | | 1.19E-02 | ↑ROR2, ↑WNT5A ...all 2 | |
| TP63 | | transcription regulator | | 1.339 | | 1.19E-02 | ↑BMP7, ↑CDK18, ...all 10 | |
| Vegf | | group | Activated | 2.007 | bias | 1.21E-02 | ↑ACKR3, ↓AN... ...all 12 | |
| HBEGF | | growth factor | | | | 1.22E-02 | ↑BCL2, ↑CXCR4,... ...all 3 | |
| U0126 | | chemical - kinase inhibi... | | -0.218 | bias | 1.24E-02 | ↑ACKR3, ↑BCL2, ...all 12 | |
| MBD3 | | enzyme | | | | 1.24E-02 | ↓FZD5, ↑WNT5A, ...all 4 | |
| STAT3 | | transcription regulator | Activated | 2.951 | bias | 1.32E-02 | ↑BCL2, ↑BMF, ↑... ...all 13 | |
| Stat5 dimer | | complex | | | | 1.35E-02 | ↑BCL2, ↓EBF1 ...all 2 | |
| EPHX2 | | enzyme | | | | 1.35E-02 | ↑HES1, ↑HEY1 ...all 2 | |
| HNRNPU | | transporter | | | | 1.35E-02 | ↓ASCL1, ↑NEUR... ...all 2 | |
| reserpine | | chemical drug | | | | 1.35E-02 | ↑ADRA2A, ↑TRH ...all 2 | |
| pilocarpine | | chemical drug | | | | 1.35E-02 | ↑NGFR, ↓NPY ...all 2 | |
| valproic acid | | chemical drug | | 0.848 | | 1.36E-02 | ↑BCL2, ↓CRH, ↑... ...all 11 | |
| quinolinic acid | | chemical - endogenous... | | | | 1.39E-02 | ↑BCL2, ↓GAD1,↑... ...all 3 | |
| K+ | | chemical - endogenous... | | | | 1.39E-02 | ↑NPTX1, ↑NR4A2, ...all 3 | |
| 4-phenylbutyric acid | | chemical - endogenous... | | 0.989 | | 1.41E-02 | ↓ABCA1, ↑BCL2,... ...all 4 | |
| NRG1 | ↓-1.933 | growth factor | | 0.684 | | 1.42E-02 | ↓ABCA1, ↓ASCL1, ...all 7 | |
| JUN | | transcription regulator | | 0.785 | bias | 1.44E-02 | ↓CRH, ↓FLNC,↑... ...all 11 | |
| BRCA1 | | transcription regulator | | -0.956 | | 1.44E-02 | ↑BCL2, ↑COL18... ...all 6 | |
| Smad2/3-Smad4 | | complex | | | | 1.51E-02 | ↑BCL2, ↑HMGA2 ...all 2 | |
| dactolisib | | chemical drug | | | | 1.51E-02 | ↑BCL2, ↓NRG1 ...all 2 | |
| quinpirole | | chemical reagent | | | | 1.51E-02 | ↓NPY, ↓SST ...all 2 | |
| NR4A1 | | ligand-dependent nucl... | | | | 1.52E-02 | ↑BCL2, ↑KCNIP3, ...all 6 | |
| STAT5B | | transcription regulator | | 0.610 | | 1.52E-02 | ↓ADGRV1, ↑BCL2, ...all 8 | |
| ADRA1B | | G-protein coupled rece... | | | | 1.56E-02 | ↑NFIX, ↓NPY, ↑... ...all 3 | |
| PRDM1 | | transcription regulator | | 1.941 | | 1.58E-02 | ↑CXCR4, ↑FNDC5 ...all 7 | |
| KMT2A | | transcription regulator | | 1.000 | bias | 1.58E-02 | ↑BCL2, ↑PRDM8, ...all 4 | |
| ITGA5 | | transmembrane receptor | | | | 1.65E-02 | ↑BCL2, ↑GAS1,↑... ...all 3 | |
| bleomycin | | chemical drug | | 1.000 | bias | 1.67E-02 | ↑ACKR3, ↑BCL2,... ...all 6 | |
| Ctbp | | group | | | | 1.69E-02 | ↑ACKR3, ↑HEY1 ...all 2 | |
| Fgfr | | group | | | | 1.69E-02 | ↑HEY1, ↑HEY2 ...all 2 | |
| RNF20 | | enzyme | | | | 1.69E-02 | ↑FEZF2, ↑NR4A2 ...all 2 | |
| ITGA6 | | transmembrane receptor | | | | 1.69E-02 | ↑BCL2, ↑ITGA2 ...all 2 | |
| ITGAL | | transmembrane receptor | | | | 1.69E-02 | ↑HES1, ↑HEY1 ...all 2 | |
| FDFT1 | | enzyme | | | | 1.69E-02 | ↓ABCA1, ↑PCSK9 ...all 2 | |
| ethosuximide | | chemical drug | | | | 1.69E-02 | ↑LEF1, ↑NEURO... ...all 2 | |
| propionic acid | | chemical - endogenous... | | | | 1.69E-02 | ↑RELN, ↓SMOC1 ...all 2 | |
| ezetimibe | | chemical drug | | | | 1.69E-02 | ↓ABCA1, ↑PCSK9 ...all 2 | |
| MYOC | | other | | | | 1.71E-02 | ↓DLX5, ↑EBF3,↑... ...all 4 | |



©2000-2018 QIAGEN. All rights reserved.

| Upstream Regul... | Expr Fold Change | Molecule Type | Predicted Activatio... | Activation z-score | Flags | p-value of ove... | Target molecule... | Mechanistic Netwo... |
|---|---|---|---|---|---|---|---|---|
| SBDS | | other | | | | 1.71E-02 | ↑CRABP1, ↑EPD... ...all 4 | |
| SOCS3 | | phosphatase | | | | 1.71E-02 | ↓ABCA1, ↑BCL2,... ...all 4 | |
| FGFR1 | | kinase | | | | 1.71E-02 | ↑BMP7, ↑HES1,... ...all 4 | |
| topotecan | | chemical drug | Inhibited | -2.449 | | 1.71E-02 | ↑DPP10, ↑ITGA2, ...all 6 | |
| HOXA3 | | transcription regulator | | | | 1.74E-02 | ↑HEY2, ↑NR2F2,... ...all 3 | |
| AIP | | transcription regulator | | | | 1.74E-02 | ↑ACKR3, ↓ADG... ...all 3 | |
| thyroid hormone | | chemical - endogenous... | | -0.256 | | 1.80E-02 | ↑NHLH1, ↓PTGDS...all 6 | |
| fluocinolone acetonide | | chemical drug | | | | 1.84E-02 | ↓ANGPTL7, ↑M... ...all 3 | |
| Hedgehog | | group | | | | 1.84E-02 | ↑GAS1, ↑HES1,... ...all 3 | |
| nitroprusside | | chemical drug | | | | 1.84E-02 | ↑BCL2, ↑LAMB1,... ...all 3 | |
| ITGB1 | | transmembrane receptor | | | | 1.84E-02 | ↑BCL2, ↑GAS1,... ...all 4 | |
| RASSF1 | | other | | 0.747 | | 1.84E-02 | ↑BCL2, ↑CXCR4,... ...all 4 | |
| KDM5B | | transcription regulator | | -0.651 | bias | 1.85E-02 | ↓ISL1, ↑NEDD9,... ...all 5 | |
| HOXA13 | | transcription regulator | | | | 1.88E-02 | ↑BMP7, ↑WNT5A ...all 2 | |
| mir-148 | | microRNA | | | | 1.88E-02 | ↓ABCA1, ↑BCL2 ...all 2 | |
| WWOX | | enzyme | | | | 1.88E-02 | ↑BCL2, ↑TP73 ...all 2 | |
| MAPK14 | | kinase | | -1.446 | bias | 1.89E-02 | ↓ABCA1, ↑BCL2,... ...all 6 | |
| H2AFB3 (includes others | | other | | | | 1.94E-02 | ↑GAS1, ↓LMO1,... ...all 3 | |
| hydrocortisone | | chemical - endogenous... | | -1.000 | | 1.96E-02 | ↓CRH, ↑CXCR4,... ...all 5 | |
| IRF8 | | transcription regulator | | 1.000 | bias | 2.01E-02 | ↑CD37, ↑IFI44L,... ...all 5 | |
| IKZF1 | | transcription regulator | Inhibited | -2.159 | | 2.07E-02 | ↓EBF1, ↑HES1,... ...all 5 | |
| MC4R | | G-protein coupled rece... | | | | 2.07E-02 | ↓NPY, ↑TRH ...all 2 | |
| mir-9 | | microRNA | | | | 2.07E-02 | ↑CXCR4, ↓GSX2 ...all 2 | |
| STK4 | | kinase | | | | 2.07E-02 | ↑BCL2, ↓WWTR1 ...all 2 | |
| plerixafor | | chemical drug | | | | 2.07E-02 | ↑CXCR4, ↓ISL1 ...all 2 | |
| finasteride | | chemical drug | | | | 2.07E-02 | ↑BCL2, ↓CRH ...all 2 | |
| Tgf beta | | group | | -0.104 | | 2.10E-02 | ↓ABCA1, ↑BCL2,... ...all 7 | |
| H89 | | chemical - kinase inhibi... | | 0.132 | bias | 2.12E-02 | ↓ABCA1, ↑BCL2,... ...all 5 | |
| bromodeoxyuridine | | chemical drug | | | | 2.14E-02 | ↑BMP7, ↑IFI44,... ...all 3 | |
| cocaine | | chemical drug | | -0.293 | bias | 2.16E-02 | ↑BCL2, ↑CACN... ...all 6 | |
| minoxidil | | chemical drug | | | | 2.22E-02 | ↑LEF1 ...all 1 | |
| amikacin | | chemical drug | | | | 2.22E-02 | ↑BCL2 ...all 1 | |
| 7beta-hydroxycholesterc | | chemical - endogenous... | | | | 2.22E-02 | ↑NR4A2 ...all 1 | |
| MICU1 | | other | | | | 2.22E-02 | ↑BCL2 ...all 1 | |
| Camkk | | group | | | | 2.22E-02 | ↓ABCA1 ...all 1 | |
| vildagliptin | | chemical drug | | | | 2.22E-02 | ↓NPY ...all 1 | |
| AEOL-10150 | | chemical drug | | | | 2.22E-02 | ↑BCL2 ...all 1 | |
| rutaecarpine | | chemical - endogenous... | | | | 2.22E-02 | ↓ABCA1 ...all 1 | |
| thymol | | chemical reagent | | | | 2.22E-02 | ↑BCL2 ...all 1 | |
| GMNC | | other | | | | 2.22E-02 | ↑FOXJ1 ...all 1 | |
| CASK | | kinase | | | | 2.22E-02 | ↑RELN ...all 1 | |
| TTC39B | | other | | | | 2.22E-02 | ↓ABCA1 ...all 1 | |
| NIF3L1 | | other | | | | 2.22E-02 | ↓ASCL1 ...all 1 | |
| GJA3 | | transporter | | | | 2.22E-02 | ↑GJA1 ...all 1 | |
| RBM15 | | other | | | | 2.22E-02 | ↑HES1 ...all 1 | |
| PCDH9 | | other | | | | 2.22E-02 | ↑BCL2 ...all 1 | |
| ISL2 | | transcription regulator | | | | 2.22E-02 | ↑ZIC2 ...all 1 | |
| hyperoside | | chemical - endogenous... | | | | 2.22E-02 | ↑BCL2 ...all 1 | |
| SENP8 | | peptidase | | | | 2.22E-02 | ↑TP73 ...all 1 | |
| EPN3 | | other | | | | 2.22E-02 | ↓EHD2 ...all 1 | |
| Pde3 | | group | | | | 2.22E-02 | ↓CRH ...all 1 | |
| ALX3 | | transcription regulator | | | | 2.22E-02 | ↓SST ...all 1 | |
| AGO3 | | translation regulator | | | | 2.22E-02 | ↑CXCR4 ...all 1 | |
| PTPase | | group | | | | 2.22E-02 | ↓ABCA1 ...all 1 | |
| CHIR-124 | | chemical reagent | | | | 2.22E-02 | ↑TP73 ...all 1 | |
| bucillamine | | chemical drug | | | | 2.22E-02 | ↑BCL2 ...all 1 | |
| MYO1C | | enzyme | | | | 2.22E-02 | ↑GSN ...all 1 | |
| CACYBP | | other | | | | 2.22E-02 | ↑BCL2 ...all 1 | |
| safflor yellow B | | chemical - endogenous... | | | | 2.22E-02 | ↑BCL2 ...all 1 | |
| NOSIP | | other | | | | 2.22E-02 | ↑BCL2 ...all 1 | |
| NPBWR1 | | G-protein coupled rece... | | | | 2.22E-02 | ↓NPY ...all 1 | |
| FANCF | | other | | | | 2.22E-02 | ↑HES1 ...all 1 | |
| SYCP3 | | other | | | | 2.22E-02 | ↑BCL2 ...all 1 | |
| mir-370 | | microRNA | | | | 2.22E-02 | ↓GAD2 ...all 1 | |
| mir-153 | | microRNA | | | | 2.22E-02 | ↑BCL2 ...all 1 | |
| miR-501-3p (and other r | | mature microRNA | | | | 2.22E-02 | ↓GAD1 ...all 1 | |
| mir-139 | | microRNA | | | | 2.22E-02 | ↑EOMES ...all 1 | |
| mir-663 | | microRNA | | | | 2.22E-02 | ↑CXCR4 ...all 1 | |
| lynestrenol | | chemical drug | | | | 2.22E-02 | ↓ABCA1 ...all 1 | |
| COPS4 | | peptidase | | | | 2.22E-02 | ↑STON2 ...all 1 | |
| CCNG1 | | other | | | | 2.22E-02 | ↑TP73 ...all 1 | |
| HIF1A-AS1 | | other | | | | 2.22E-02 | ↑BCL2 ...all 1 | |
| PIM | | group | | | | 2.22E-02 | ↑BCL2 ...all 1 | |
| CRHBP | | other | | | | 2.22E-02 | ↓CRH ...all 1 | |
| RPL15 | | other | | | | 2.22E-02 | ↑HMGA2 ...all 1 | |
| PLIN3 | | other | | | | 2.22E-02 | ↑BCL2 ...all 1 | |
| cyclo(iso-Asp-GR)-LLIIKL | | chemical reagent | | | | 2.22E-02 | ↑BCL2 ...all 1 | |
| MAML3 | | transcription regulator | | | | 2.22E-02 | ↑HES1 ...all 1 | |
| [illegible] | | translation regulator | | | | 2.22E-02 | ↑CXCR4 ...all 1 | |



©2000-2018 QIAGEN. All rights reserved.

| Upstream Regul... | Expr Fold Change | Molecule Type | Predicted Activatio... | Activation z-score | Flags | p-value of ove... | Target molecule... | Mechanistic Netwo... |
|---|---|---|---|---|---|---|---|---|
| AGO4 | | translation regulator | | | | 2.22E-02 | ↑CXCR4 ...all 1 | |
| CREB5 | | transcription regulator | | | | 2.22E-02 | ↓CRH ...all 1 | |
| NPAS3 | | transcription regulator | | | | 2.22E-02 | ↑RELN ...all 1 | |
| LRP1B | | transmembrane receptor | | | | 2.22E-02 | ↓ABCA1 ...all 1 | |
| TH | | enzyme | | | | 2.22E-02 | ↓NPY ...all 1 | |
| FOXL1 | | transcription regulator | | | | 2.22E-02 | ↑WNT5A ...all 1 | |
| STAMBP | | enzyme | | | | 2.22E-02 | ↑CXCR4 ...all 1 | |
| PREB | | transcription regulator | | | | 2.22E-02 | ↓ABCA1 ...all 1 | |
| N-[4-chloro-3-(trifluoror | | chemical reagent | | | | 2.22E-02 | ↑NEUROD1 ...all 1 | |
| TRIM54 | | other | | | | 2.22E-02 | ↓FLNC ...all 1 | |
| NEU2 | | enzyme | | | | 2.22E-02 | ↑BCL2 ...all 1 | |
| UBE2V1 | | transcription regulator | | | | 2.22E-02 | ↑BCL2 ...all 1 | |
| Gm4836 (includes others | | other | | | | 2.22E-02 | ↑BCL2 ...all 1 | |
| AP2B1 | | transporter | | | | 2.22E-02 | ↓PTGDS ...all 1 | |
| MAML2 | | transcription regulator | | | | 2.22E-02 | ↑HES1 ...all 1 | |
| CLEC2D | | transmembrane receptor | | | | 2.22E-02 | ↑CXCR4 ...all 1 | |
| VNN1 | | enzyme | | | | 2.22E-02 | ↑BCL2 ...all 1 | |
| LAMB2 | | enzyme | | | | 2.22E-02 | ↑LAMB1 ...all 1 | |
| 1,5-bis-(dihexyl-N-nitros | | chemical reagent | | | | 2.22E-02 | ↓PDE5A ...all 1 | |
| 3830403N18Rik/XI | | other | | | | 2.22E-02 | ↑BCL2 ...all 1 | |
| securinine | | chemical reagent | | | | 2.22E-02 | ↑TP73 ...all 1 | |
| 1,1-bis(3'-indolyl)-1-(4-h | | chemical reagent | | | | 2.22E-02 | ↑BCL2 ...all 1 | |
| S7 | | chemical - kinase inhibi... | | | | 2.22E-02 | ↑BCL2 ...all 1 | |
| SI163 | | chemical - kinase inhibi... | | | | 2.22E-02 | ↑BCL2 ...all 1 | |
| S29 | | chemical - kinase inhibi... | | | | 2.22E-02 | ↑BCL2 ...all 1 | |
| daphnoretin | | chemical - endogenous... | | | | 2.22E-02 | ↑BCL2 ...all 1 | |
| methoxamine | | chemical drug | | | | 2.22E-02 | ↓CRH ...all 1 | |
| PSB-1115 | | chemical reagent | | | | 2.22E-02 | ↓PDE5A ...all 1 | |
| XK469 | | chemical drug | | | | 2.22E-02 | ↑BCL2 ...all 1 | |
| talipexole | | chemical drug | | | | 2.22E-02 | ↑BCL2 ...all 1 | |
| gabazine | | chemical drug | | | | 2.22E-02 | ↑GJA1 ...all 1 | |
| bumetanide | | chemical drug | | | | 2.22E-02 | ↑NGFR ...all 1 | |
| hexaarginine-neomycin | | chemical reagent | | | | 2.22E-02 | ↑CXCR4 ...all 1 | |
| AN-207 | | chemical toxicant | | | | 2.22E-02 | ↑BCL2 ...all 1 | |
| SSR18057 | | chemical drug | | | | 2.22E-02 | ↑BCL2 ...all 1 | |
| NSC651016 | | chemical reagent | | | | 2.22E-02 | ↑CXCR4 ...all 1 | |
| fenoprofen | | chemical drug | | | | 2.22E-02 | ↑NGFR ...all 1 | |
| calcium-EDTA | | chemical drug | | | | 2.22E-02 | ↑NGFR ...all 1 | |
| isoalantolactone | | chemical - endogenous... | | | | 2.22E-02 | ↑BCL2 ...all 1 | |
| swainsonine | | chemical - endogenous... | | | | 2.22E-02 | ↑BCL2 ...all 1 | |
| ULK4 | | kinase | | | | 2.27E-02 | ↑FOXJ1, ↑GSN ...all 2 | |
| H2AFZ | | other | | | | 2.27E-02 | ↑CXCR4, ↑TTR ...all 2 | |
| miR-103-3p (and other r | | mature microRNA | | | | 2.27E-02 | ↑CACNA2D1, ↑... ...all 2 | |
| PCDH11Y | | other | | | | 2.27E-02 | ↑GJA1, ↑WNT7B ...all 2 | |
| olanzapine | | chemical drug | | 1.964 | | 2.35E-02 | ↑BCL2, ↑NR4A2,... ...all 4 | |
| dihydrotestosterone | | chemical - endogenous... | | -0.503 | bias | 2.36E-02 | ↓ABCA1, ↑ACK... ...all 12 | |
| leukotriene D4 | | chemical - endogenous... | | | | 2.36E-02 | ↑KCNJ2, ↑NEDD9,...all 3 | |
| oblimersen | | biologic drug | | | | 2.36E-02 | ↑BCL2, ↑IFI44, ↑... ...all 3 | |
| anisomycin | | chemical - endogenous... | | | | 2.36E-02 | ↑BCL2, ↑GJA1, ↑... ...all 3 | |
| ERG | | transcription regulator | | | | 2.42E-02 | ↑CXCR4, ↓FLNC,... ...all 6 | |
| NCOA1 | | transcription regulator | | -0.152 | | 2.43E-02 | ↓ABCA1, ↓CRH,... ...all 5 | |
| aspirin | | chemical drug | | -1.225 | bias | 2.43E-02 | ↓ABCA1, ↑BCL2,... ...all 4 | |
| EDN1 | | cytokine | | 0.933 | bias | 2.47E-02 | ↑BCL2, ↑CXCR4,... ...all 6 | |
| NCOR2 | | transcription regulator | | | | 2.48E-02 | ↑BCL2, ↓CRH, ↑... ...all 3 | |
| FOXC2 | | transcription regulator | | | | 2.48E-02 | ↑CXCR4, ↑LEF1,... ...all 3 | |
| SP1 | | transcription regulator | | 1.842 | bias | 2.48E-02 | ↓ABCA1, ↑BCL2, ...all 12 | |
| prostaglandin A1 | | chemical - endogenous... | | | | 2.48E-02 | ↑BCL2, ↓GAD1 ...all 2 | |
| Pkg | | group | | | | 2.48E-02 | ↑BCL2, ↑LEF1 ...all 2 | |
| ADCY | | group | | | | 2.48E-02 | ↑GJA1, ↑NR4A2 ...all 2 | |
| IKK (complex) | | complex | | | | 2.48E-02 | ↑BCL2, ↑EOMES ...all 2 | |
| OXT | | other | | | | 2.48E-02 | ↓CRH, ↑GJA1 ...all 2 | |
| mir-124 | | microRNA | | | | 2.48E-02 | ↓DLX5, ↑LHX2 ...all 2 | |
| KISS1 | | other | | | | 2.48E-02 | ↓NPY, ↑OTX2 ...all 2 | |
| NOV | | growth factor | | | | 2.48E-02 | ↑HES1, ↑ITGA2 ...all 2 | |
| CRHR2 | | G-protein coupled rece... | | | | 2.48E-02 | ↑BCL2, ↓CRH ...all 2 | |
| LIPG | | enzyme | | | | 2.48E-02 | ↓ABCA1, ↑BCL2 ...all 2 | |
| IL15RA | | transmembrane receptor | | | | 2.48E-02 | ↑BCL2, ↓GAD2 ...all 2 | |
| PD18435 | | chemical drug | | | | 2.48E-02 | ↑BCL2, ↑TP73 ...all 2 | |
| ketamine | | chemical drug | | | | 2.48E-02 | ↓GAD1, ↑NEUR... ...all 2 | |
| wogonin | | chemical - endogenous... | | | | 2.48E-02 | ↓ABCA1, ↑BCL2 ...all 2 | |
| HNRNPA2B1 | | other | | | | 2.49E-02 | ↑GJA1, ↑GSN, ↑... ...all 5 | |
| geldanamycin | | chemical - endogenous... | | -0.447 | bias | 2.49E-02 | ↑BCL2, ↑GJA1, ↑... ...all 5 | |
| bucladesine | | chemical toxicant | | -0.022 | bias | 2.62E-02 | ↓ABCA1, ↑BCL2,... ...all 8 | |
| HMGA1 | | transcription regulator | | 0.651 | | 2.69E-02 | ↑BCL2, ↑GSN, ↑... ...all 5 | |
| mir-33 | | microRNA | | | | 2.70E-02 | ↓ABCA1, ↑HMG... ...all 2 | |
| ETV1 | | transcription regulator | | | | 2.70E-02 | ↓ASCL1, ↑CXCR4 ...all 2 | |
| HSPA9 | | other | | | | 2.70E-02 | ↑HES1, ↑NEDD9 ...all 2 | |
| pitavastatin | | chemical drug | | | | 2.70E-02 | ↓ABCA1, ↑PCSK9 ...all 2 | |
| desipramine | | chemical drug | | | | 2.70E-02 | ↑ADRA2A, ↑BCL2 ...all 2 | |
| [illegible] | | [illegible] | | | | [illegible] | [illegible] | |



©2000-2018 QIAGEN. All rights reserved.

| Upstream Regul... | Expr Fold Change | Molecule Type | Predicted Activatio... | Activation z-score | Flags | p-value of ove... | Target molecule... | Mechanistic Netwo... |
|---|---|---|---|---|---|---|---|---|
| CSHL1 | | growth factor | | | | 2.83E-02 | ↓NPY, ↓SST, ↓T... ...all 3 | |
| pravastatin | | chemical drug | | | | 2.83E-02 | ↓ABCA1, ↑GJA1, ... ...all 3 | |
| GFI1 | | transcription regulator | | | | 2.85E-02 | ↑CXCR4, ↑GJA1, ... ...all 4 | |
| Ikb | | group | | | | 2.92E-02 | ↑BCL2, ↑TP73 ...all 2 | |
| TRPS1 | | transcription regulator | | | | 2.92E-02 | ↑SEMA6A, ↑SER... ...all 2 | |
| GRHL2 | | transcription regulator | | | | 2.92E-02 | ↑BCL2, ↓SEMA3C ...all 2 | |
| miR-181a-5p (and other | | mature microRNA | | | | 2.92E-02 | ↑BCL2, ↓VSNL1 ...all 2 | |
| AKT3 | | kinase | | | | 2.92E-02 | ↑BMF, ↑RELN ...all 2 | |
| SIX5 | | transcription regulator | | | | 2.92E-02 | ↑COL9A2, ↑EBF2 ...all 2 | |
| GNL3 | | other | | | | 2.92E-02 | ↑EOMES, ↑MSX1 ...all 2 | |
| Cdkn1c | | other | | | | 2.92E-02 | ↑CXCR4, ↑NGFR ...all 2 | |
| NFATC1 | | transcription regulator | | | | 2.94E-02 | ↑BCL2, ↑BMP7, ... ...all 4 | |
| DDIT3 | | transcription regulator | | | | 2.96E-02 | ↑BCL2, ↑BHLHE... ...all 3 | |
| RNF2 | | transcription regulator | | | | 2.96E-02 | ↑EOMES, ↑HEY2, ...all 3 | |
| CHD4 | | enzyme | | | | 2.96E-02 | ↑NGFR, ↑NHLH1, ...all 3 | |
| CTGF | | growth factor | | 1.109 | bias | 3.03E-02 | ↑BCL2, ↑HES1, ... ...all 4 | |
| BMP | | group | | | | 3.15E-02 | ↓ISL1, ↑MSX1 ...all 2 | |
| SSTR2 | | G-protein coupled rece... | | | | 3.15E-02 | ↑BCL2, ↓SST ...all 2 | |
| BAG1 | | other | | | | 3.15E-02 | ↑BCL2, ↑TP73 ...all 2 | |
| UCN-01 | | chemical drug | | | | 3.15E-02 | ↑BCL2, ↑TP73 ...all 2 | |
| LY294002 | | chemical - kinase inhibi... | | -1.598 | bias | 3.19E-02 | ↓ABCA1, ↑ACK... ...all 11 | |
| NFYB | | transcription regulator | | | | 3.20E-02 | ↑BCL2, ↓CACN... ...all 7 | |
| mir-34 | | microRNA | | | | 3.22E-02 | ↑BCL2, ↑HEY1, ... ...all 3 | |
| Sn50 peptide | | chemical toxicant | | | | 3.22E-02 | ↑BCL2, ↑GJA1, ↓... ...all 3 | |
| CHUK | | kinase | | 1.633 | bias | 3.23E-02 | ↑ACKR3, ↑COL1... ...all 6 | |
| SIM1 | | transcription regulator | | -0.128 | bias | 3.29E-02 | ↓CRH, ↓POU3F2, ...all 6 | |
| RAF1 | | kinase | | 0.355 | bias | 3.29E-02 | ↑BCL2, ↑CD37, ... ...all 6 | |
| simvastatin | | chemical drug | | -0.483 | | 3.29E-02 | ↓ABCA1, ↓ASL, ... ...all 6 | |
| propylthiouracil | | chemical drug | | | | 3.31E-02 | ↑GSN, ↓ROBO1, ... ...all 4 | |
| L-tyrosine | | chemical - endogenous... | | | | 3.32E-02 | ↑BCL2 ...all 1 | |
| pentosan polysulfate | | chemical drug | | | | 3.32E-02 | ↓ABCA1 ...all 1 | |
| stigmasterol | | chemical - endogenous... | | | | 3.32E-02 | ↓ABCA1 ...all 1 | |
| hyodeoxycholic acid | | chemical - endogenous... | | | | 3.32E-02 | ↓ABCA1 ...all 1 | |
| DUB | | group | | | | 3.32E-02 | ↓ABCA1 ...all 1 | |
| PLC gamma | | group | | | | 3.32E-02 | ↓SCN9A ...all 1 | |
| sethoxydim | | chemical toxicant | | | | 3.32E-02 | ↑BCL2 ...all 1 | |
| FREM1 | ↑2.552 | other | | | | 3.32E-02 | ↑FREM2 ...all 1 | |
| SLC6A12 | | transporter | | | | 3.32E-02 | ↑NEUROD1 ...all 1 | |
| SPRED2 | | cytokine | | | | 3.32E-02 | ↓CRH ...all 1 | |
| C1QL1 | | other | | | | 3.32E-02 | ↓CRH ...all 1 | |
| LAPTM4B | | other | | | | 3.32E-02 | ↑BCL2 ...all 1 | |
| Integrin alpha 4 beta 1 | | complex | | | | 3.32E-02 | ↑BCL2 ...all 1 | |
| CKLF | | cytokine | | | | 3.32E-02 | ↑BCL2 ...all 1 | |
| xenon | | chemical drug | | | | 3.32E-02 | ↑BCL2 ...all 1 | |
| LHX8 | | transcription regulator | | | | 3.32E-02 | ↓ISL1 ...all 1 | |
| ACAT1 | | enzyme | | | | 3.32E-02 | ↓ABCA1 ...all 1 | |
| GEM231 | | chemical drug | | | | 3.32E-02 | ↑BCL2 ...all 1 | |
| cerebrolysin | | chemical drug | | | | 3.32E-02 | ↑BCL2 ...all 1 | |
| RNF144B | | enzyme | | | | 3.32E-02 | ↑TP73 ...all 1 | |
| PLEC | | other | | | | 3.32E-02 | ↑CXCR4 ...all 1 | |
| 17alpha-hydroxyprogest | | chemical drug | | | | 3.32E-02 | ↓CRH ...all 1 | |
| miR-136-5p (miRNAs w/ | | mature microRNA | | | | 3.32E-02 | ↑BCL2 ...all 1 | |
| miR-370-3p (and other n | | mature microRNA | | | | 3.32E-02 | ↓GAD2 ...all 1 | |
| miR-153-3p (miRNAs w/ | | mature microRNA | | | | 3.32E-02 | ↑BCL2 ...all 1 | |
| HOXB2 | | transcription regulator | | | | 3.32E-02 | ↑OTX2 ...all 1 | |
| MCHR1 | | G-protein coupled rece... | | | | 3.32E-02 | ↑TRH ...all 1 | |
| TLE4 | | transcription regulator | | | | 3.32E-02 | ↓SIX3 ...all 1 | |
| COL7A1 | | other | | | | 3.32E-02 | ↑WNT5A ...all 1 | |
| NRP2 | | kinase | | | | 3.32E-02 | ↑BCL2 ...all 1 | |
| N4BP1 | | other | | | | 3.32E-02 | ↑TP73 ...all 1 | |
| ANKRD1 | | transcription regulator | | | | 3.32E-02 | ↑BCL2 ...all 1 | |
| LSS | | enzyme | | | | 3.32E-02 | ↓ABCA1 ...all 1 | |
| ZNF350 | | transcription regulator | | | | 3.32E-02 | ↑HMGA2 ...all 1 | |
| TOP2B | | enzyme | | | | 3.32E-02 | ↑RELN ...all 1 | |
| SPEN | | transcription regulator | | | | 3.32E-02 | ↑HES1 ...all 1 | |
| HELT | | transcription regulator | | | | 3.32E-02 | ↑NEUROG2 ...all 1 | |
| CCNL2 | | other | | | | 3.32E-02 | ↑BCL2 ...all 1 | |
| MAP1S | | enzyme | | | | 3.32E-02 | ↑BCL2 ...all 1 | |
| AP2A2 | | transporter | | | | 3.32E-02 | ↑BCL2 ...all 1 | |
| RBBP8 | | enzyme | | | | 3.32E-02 | ↑HMGA2 ...all 1 | |
| MATK | | kinase | | | | 3.32E-02 | ↑CXCR4 ...all 1 | |
| ARHGEF11 | | other | | | | 3.32E-02 | ↓ABCA1 ...all 1 | |
| GCM1 | | transcription regulator | | | | 3.32E-02 | ↓FZD5 ...all 1 | |
| MT3 | | other | | | | 3.32E-02 | ↑BCL2 ...all 1 | |
| CNTFR | | transmembrane receptor | | | | 3.32E-02 | ↑GJA1 ...all 1 | |
| WTAP | | other | | | | 3.32E-02 | ↑BCL2 ...all 1 | |
| cilomilast | | chemical drug | | | | 3.32E-02 | ↑CXCR4 ...all 1 | |
| ethyl ether | | chemical reagent | | | | 3.32E-02 | ↓CRH ...all 1 | |
| tarenflurbil | | chemical drug | | | | 3.32E-02 | ↑NGFR ...all 1 | |



©2000-2018 QIAGEN. All rights reserved.

| Upstream Regul... | Expr Fold Change | Molecule Type | Predicted Activatio... | Activation z-score | Flags | p-value of ove... | Target molecule... | Mechanistic Netwo... |
|---|---|---|---|---|---|---|---|---|
| minodronate | | chemical drug | | | | 3.32E-02 | ↑CXCR4 ...all 1 | |
| pramipexole | | chemical drug | | | | 3.32E-02 | ↑BCL2 ...all 1 | |
| lead nitrate | | chemical toxicant | | | | 3.32E-02 | ↑BCL2 ...all 1 | |
| imisopasem manganese | | chemical drug | | | | 3.32E-02 | ↑BCL2 ...all 1 | |
| (Ala9)-autocamtide-2 | | chemical - kinase inhibi... | | | | 3.32E-02 | ↓CRH ...all 1 | |
| atosiban | | biologic drug | | | | 3.32E-02 | ↑GJA1 ...all 1 | |
| lisofylline | | chemical drug | | | | 3.32E-02 | ↑LAMB1 ...all 1 | |
| 10-hydroxycamptothecir | | chemical - endogenous... | | | | 3.32E-02 | ↑BCL2 ...all 1 | |
| 3,3',5-triiodothyroacetic | | chemical - endogenous... | | | | 3.32E-02 | ↑TRH ...all 1 | |
| 2R,4R-4-aminopyrrolidin | | chemical reagent | | | | 3.32E-02 | ↑TP73 ...all 1 | |
| lipopolysaccharide | | chemical drug | Activated | 2.836 | bias | 3.35E-02 | ↓ABCA1, ↑ACK... ...all 28 | |
| gemcitabine | | chemical drug | | | | 3.35E-02 | ↑BCL2, ↑CXCR4, ... ...all 3 | |
| LIF | | cytokine | | | | 3.35E-02 | ↑BMP7, ↑HES1, ... ...all 6 | |
| MAC | | complex | | | | 3.39E-02 | ↑BCL2, ↑RGS16 ...all 2 | |
| Pka catalytic subunit | | group | | | | 3.39E-02 | ↓ABCA1, ↓SST ...all 2 | |
| CD9 | | other | | | | 3.39E-02 | ↑ITGA2, ↑WNT5A ...all 2 | |
| nitric oxide | | chemical - endogenous... | | | | 3.41E-02 | ↑BCL2, ↓CRH, ↑... ...all 5 | |
| PTH | | other | | 1.961 | bias | 3.49E-02 | ↑BCL2, ↑CXCR4, ... ...all 5 | |
| NRG2 | | growth factor | | | | 3.63E-02 | ↑BCL2, ↓GAD1, ... ...all 3 | |
| SMARCA2 | | transcription regulator | | | | 3.63E-02 | ↑BCL2, ↑HES1, ... ...all 3 | |
| OTX2 | ↑1.657 | transcription regulator | | | | 3.63E-02 | ↓SEMA3C, ↓SIX3, ...all 3 | |
| entinostat | | chemical drug | | | | 3.63E-02 | ↑BCL2, ↑GSN, ↑... ...all 3 | |
| NTF3 | | growth factor | | | | 3.63E-02 | ↑GJA1, ↓SST ...all 2 | |
| mir-22 | | microRNA | | | | 3.63E-02 | ↑BMP7, ↑LEF1 ...all 2 | |
| USP7 | | peptidase | | | | 3.63E-02 | ↑ACKR3, ↑BMF ...all 2 | |
| ASAH1 | | enzyme | | | | 3.63E-02 | ↑NR4A2, ↑TSPO ...all 2 | |
| CPE | | peptidase | | | | 3.63E-02 | ↑BCL2, ↑TRH ...all 2 | |
| Pkc(s) | | group | | -0.065 | bias | 3.69E-02 | ↓ABCA1, ↑BCL2, ... ...all 6 | |
| PI3K (family) | | group | | 0.940 | bias | 3.71E-02 | ↓ABCA1, ↑BCL2, ... ...all 4 | |
| TBX5 | | transcription regulator | | | | 3.77E-02 | ↑GJA1, ↑HEY2, ↑... ...all 3 | |
| ESR1 | | ligand-dependent nucl... | | 0.362 | | 3.78E-02 | ↑ABCA4, ↑ACK... ...all 22 | |
| HMGN1 | | transcription regulator | | | | 3.89E-02 | ↑NEUROD1, ↑TTR ...all 2 | |
| letrozole | | chemical drug | | | | 3.89E-02 | ↑BCL2, ↑GJA1 ...all 2 | |
| apomorphine | | chemical drug | | | | 3.89E-02 | ↑BCL2, ↑GJA1 ...all 2 | |
| hyaluronic acid | | chemical - endogenous... | | | | 3.92E-02 | ↑ACKR3, ↑BCL2, ... ...all 4 | |
| HMOX1 | | enzyme | | | | 3.92E-02 | ↓ABCA1, ↑BCL2, ... ...all 4 | |
| RET | | kinase | | | | 3.92E-02 | ↑BCL2, ↑CXCR4, ... ...all 4 | |
| IL1 | | group | | 0.617 | bias | 3.96E-02 | ↑BCL2, ↓CRH, ↓... ...all 8 | |
| NR3C1 | | ligand-dependent nucl... | | 1.109 | | 4.00E-02 | ↓ABCA1, ↓AN... ...all 13 | |
| estradiol benzoate | | chemical drug | | | | 4.06E-02 | ↑GJA1, ↓NPY, ↓... ...all 3 | |
| Raf | | group | | | | 4.06E-02 | ↑HMGA2, ↓SEM... ...all 3 | |
| GHRL | | growth factor | | | | 4.06E-02 | ↑BCL2, ↓NPY, ↓... ...all 3 | |
| MAPK8 | | kinase | | | | 4.07E-02 | ↓ABCA1, ↑BCL2, ... ...all 5 | |
| L-arginine | | chemical - endogenous... | | | | 4.14E-02 | ↓ASL, ↑BCL2 ...all 2 | |
| SMTNL1 | | other | | | | 4.14E-02 | ↓FLNC, ↑GJA1 ...all 2 | |
| CRTC2 | | other | | | | 4.14E-02 | ↑BCL2, ↓CRH ...all 2 | |
| ARRB1 | | other | | | | 4.14E-02 | ↑BCL2, ↑LEF1 ...all 2 | |
| androgen | | chemical drug | | | | 4.16E-02 | ↑BCL2, ↑CXCR4, ... ...all 5 | |
| ANXA7 | | ion channel | | | | 4.21E-02 | ↑ITGA2, ↑NEUR... ...all 3 | |
| COLQ | | other | | | | 4.21E-02 | ↑ADAMTS2, ↑L... ...all 3 | |
| lipoteichoic acid | | chemical - endogenous... | | | | 4.21E-02 | ↑BCL2, ↑CXCR4, ... ...all 3 | |
| trans-hydroxytamoxifen | | chemical drug | | -1.982 | | 4.24E-02 | ↑COL4A5, ↑LA... ...all 4 | |
| miR-199a-5p (and other | | mature microRNA | | | | 4.37E-02 | ↑COL4A5, ↑NED... ...all 3 | |
| 10E,12Z-octadecadienoi | | chemical - endogenous... | | | | 4.37E-02 | ↑BCL2, ↓EBF1, ↑... ...all 3 | |
| 1-O-hexadecyl-2-N-met | | chemical - endogenous... | | | | 4.40E-02 | ↓SST ...all 1 | |
| LRP4 | | other | | | | 4.40E-02 | ↑BMP7 ...all 1 | |
| OSGIN1 | | growth factor | | | | 4.40E-02 | ↑BCL2 ...all 1 | |
| SPRED1 | | other | | | | 4.40E-02 | ↓CRH ...all 1 | |
| PROM1 | | other | | | | 4.40E-02 | ↑NR4A2 ...all 1 | |
| OLFM2 | | other | | | | 4.40E-02 | ↑HEY2 ...all 1 | |
| Meg3 | | other | | | | 4.40E-02 | ↑HES1 ...all 1 | |
| ATOH7 | | other | | | | 4.40E-02 | ↑BARHL2 ...all 1 | |
| CRTAM | | other | | | | 4.40E-02 | ↑EOMES ...all 1 | |
| enecadin | | chemical drug | | | | 4.40E-02 | ↑BCL2 ...all 1 | |
| C1GALT1 | | enzyme | | | | 4.40E-02 | ↑BCL2 ...all 1 | |
| HMX2 | | transcription regulator | | | | 4.40E-02 | ↓DLX5 ...all 1 | |
| WNT9B | | other | | | | 4.40E-02 | ↑LHX1 ...all 1 | |
| MITF-p300/CBP | | complex | | | | 4.40E-02 | ↑BCL2 ...all 1 | |
| HHIP | | other | | | | 4.40E-02 | ↓EBF1 ...all 1 | |
| firtecan pegol | | chemical drug | | | | 4.40E-02 | ↑CXCR4 ...all 1 | |
| KCNIP1 | | ion channel | | | | 4.40E-02 | ↑DPP10 ...all 1 | |
| CTSD | | peptidase | | | | 4.40E-02 | ↓ABCA1 ...all 1 | |
| RNF6 | | transcription regulator | | | | 4.40E-02 | ↑BMF ...all 1 | |
| PAK4 | | kinase | | | | 4.40E-02 | ↑BCL2 ...all 1 | |
| KCND3 | | ion channel | | | | 4.40E-02 | ↑DPP10 ...all 1 | |
| MAL | | other | | | | 4.40E-02 | ↑NGFR ...all 1 | |
| RPS11 | | other | | | | 4.40E-02 | ↑HMGA2 ...all 1 | |
| ARTN | | growth factor | | | | 4.40E-02 | ↑BCL2 ...all 1 | |
| MS4A2 | | transmembrane receptor | | | | 4.40E-02 | ↑BCL2 ...all 1 | |



©2000-2018 QIAGEN. All rights reserved.

| Upstream Regul... | Expr Fold Change | Molecule Type | Predicted Activatio... | Activation z-score | Flags | p-value of ove... | Target molecule... | Mechanistic Netwo... |
|---|---|---|---|---|---|---|---|---|
| PHF19 | | other | | | | 4.40E-02 | ↑OTX2 ...all 1 | |
| miR-515-5p (and other r | | mature microRNA | | | | 4.40E-02 | ↑TCF7L1 ...all 1 | |
| miR-516a-3p (and other | | mature microRNA | | | | 4.40E-02 | ↑WNT5A ...all 1 | |
| miR-202-3p (and other n | | mature microRNA | | | | 4.40E-02 | ↑LRIG3 ...all 1 | |
| miR-193a-5p (miRNAs w | | mature microRNA | | | | 4.40E-02 | ↑TP73 ...all 1 | |
| LPCAT3 | | enzyme | | | | 4.40E-02 | ↓ABCA1 ...all 1 | |
| CBFA2T2 | | transcription regulator | | | | 4.40E-02 | ↑HES1 ...all 1 | |
| NUCB2 | | other | | | | 4.40E-02 | ↑TRH ...all 1 | |
| ANKH | | transporter | | | | 4.40E-02 | ↑WNT5A ...all 1 | |
| TIAM1 | | other | | | | 4.40E-02 | ↑LEF1 ...all 1 | |
| PHLDA1 | | other | | | | 4.40E-02 | ↑ITGA2 ...all 1 | |
| CYP8B1 | | enzyme | | | | 4.40E-02 | ↓ABCA1 ...all 1 | |
| HSPB6 | | other | | | | 4.40E-02 | ↑BCL2 ...all 1 | |
| SIGMAR1 | | transmembrane receptor | | | | 4.40E-02 | ↑BCL2 ...all 1 | |
| DCC | | transmembrane receptor | | | | 4.40E-02 | ↑TP73 ...all 1 | |
| CELF2 | | other | | | | 4.40E-02 | ↑BCL2 ...all 1 | |
| FOXE1 | | transcription regulator | | | | 4.40E-02 | ↑WNT5A ...all 1 | |
| AES | | transcription regulator | | | | 4.40E-02 | ↓SIX3 ...all 1 | |
| 2-[[4-[(e)-styryl]phenoxy | | chemical reagent | | | | 4.40E-02 | ↓ABCA1 ...all 1 | |
| LGALS2 | | other | | | | 4.40E-02 | ↑BCL2 ...all 1 | |
| RPS18 | | other | | | | 4.40E-02 | ↑HMGA2 ...all 1 | |
| CSNK1A1 | | kinase | | | | 4.40E-02 | ↑CXCR4 ...all 1 | |
| TSN | | other | | | | 4.40E-02 | ↑BCL2 ...all 1 | |
| SCIN | | other | | | | 4.40E-02 | ↑GSN ...all 1 | |
| RPL35A | | other | | | | 4.40E-02 | ↑HMGA2 ...all 1 | |
| RPL12 | | other | | | | 4.40E-02 | ↑HMGA2 ...all 1 | |
| RPL5 | | other | | | | 4.40E-02 | ↑HMGA2 ...all 1 | |
| griseofulvin | | chemical drug | | | | 4.40E-02 | ↑GJA1 ...all 1 | |
| neopterin | | chemical - endogenous... | | | | 4.40E-02 | ↓ABCA1 ...all 1 | |
| 1-butanol | | chemical - endogenous... | | | | 4.40E-02 | ↓ABCA1 ...all 1 | |
| mianserin | | chemical drug | | | | 4.40E-02 | ↓CRH ...all 1 | |
| NSC719239 | | chemical drug | | | | 4.40E-02 | ↑ITGA2 ...all 1 | |
| TAS-103 | | chemical drug | | | | 4.40E-02 | ↑BCL2 ...all 1 | |
| pyridoxal phosphate | | chemical - endogenous... | | | | 4.40E-02 | ↑BCL2 ...all 1 | |
| ibotenic acid | | chemical toxicant | | | | 4.40E-02 | ↓CRH ...all 1 | |
| sodium chlorate | | chemical reagent | | | | 4.40E-02 | ↑BMP7 ...all 1 | |
| chlorophyll a | | chemical - endogenous... | | | | 4.40E-02 | ↑BCL2 ...all 1 | |
| retinaldehyde | | chemical - endogenous... | | | | 4.40E-02 | ↓PTGDS ...all 1 | |
| insulin glargine | | chemical drug | | | | 4.40E-02 | ↓NPY ...all 1 | |
| limonene | | chemical - endogenous... | | | | 4.40E-02 | ↓NPY ...all 1 | |
| emetine | | chemical toxicant | | | | 4.40E-02 | ↑SLC17A7 ...all 1 | |
| galanthamine | | chemical drug | | | | 4.40E-02 | ↑BCL2 ...all 1 | |
| vigabatrin | | chemical drug | | | | 4.40E-02 | ↓GAD1 ...all 1 | |
| steroid hormone | | chemical - other | | | | 4.40E-02 | ↑CXCR4 ...all 1 | |
| S-nitroso-N-acetyl-DL-p | | chemical reagent | | | | 4.41E-02 | ↑BCL2, ↑TP73 ...all 2 | |
| TCF7L1 | ↑1.657 | transcription regulator | | | | 4.41E-02 | ↑EOMES, ↑TNF... ...all 2 | |
| Focal adhesion kinase | | group | | | | 4.41E-02 | ↑BCL2, ↑NEURO... ...all 2 | |
| WNT11 | | other | | | | 4.41E-02 | ↓ASCL1, ↑RSPO2 ...all 2 | |
| betulinic acid | | chemical drug | | | | 4.41E-02 | ↓ABCA1, ↑BCL2 ...all 2 | |
| miR-124-3p (and other r | | mature microRNA | | -1.039 | bias | 4.42E-02 | ↓DLX2, ↑GSN, ↑... ...all 6 | |
| Growth hormone | | group | | -0.447 | | 4.50E-02 | ↑BCL2, ↑BMP7, ↑... ...all 6 | |
| testosterone | | chemical - endogenous... | | -1.267 | bias | 4.62E-02 | ↑ADRA2A, ↑BCL2, ...all 8 | |
| WNT4 | | cytokine | | | | 4.68E-02 | ↑GAS1, ↑LHX1 ...all 2 | |
| CNOT7 | | transcription regulator | | | | 4.68E-02 | ↑IFI44L, ↓PPM1K ...all 2 | |
| DIO3 | | enzyme | | | | 4.68E-02 | ↑NEUROD1, ↑T... ...all 2 | |
| BECN1 | | other | | | | 4.68E-02 | ↑BCL2, ↑BMF ...all 2 | |
| DRD2 | | G-protein coupled rece... | | | | 4.68E-02 | ↓GAD1, ↑KCNJ2 ...all 2 | |
| PBX1 | | transcription regulator | | | | 4.68E-02 | ↓ISL1, ↓SST ...all 2 | |
| fluvoxamine | | chemical drug | | | | 4.68E-02 | ↑CD37, ↑NR4A2 ...all 2 | |
| mir-29 | | microRNA | | | | 4.68E-02 | ↑ADAMTS9, ↑B... ...all 3 | |
| GCG | | other | | | | 4.68E-02 | ↑ACKR3, ↑BCL2, ...all 3 | |
| MEF2C | | transcription regulator | | | | 4.70E-02 | ↑CXCR4, ↑GJA1, ...all 4 | |
| SNAI1 | | transcription regulator | | | | 4.70E-02 | ↓FLNC, ↑GSN, ↑... ...all 4 | |
| PI3K (complex) | | complex | | 1.000 | bias | 4.78E-02 | ↓ABCA1, ↑BCL2, ...all 7 | |
| HOXA9 | | transcription regulator | | | | 4.80E-02 | ↑BCL2, ↓FZD5, ↑... ...all 5 | |
| NTRK2 | | kinase | | | | 4.85E-02 | ↑AMBN, ↓GAD1, ...all 3 | |
| NFKB2 | | transcription regulator | | | | 4.85E-02 | ↑BCL2, ↓CRH, ↑... ...all 3 | |
| PAX7 | | transcription regulator | | | | 4.85E-02 | ↓ABCA1, ↑CXCR4 ...all 3 | |
| IL17A | | cytokine | | -0.730 | bias | 4.90E-02 | ↑BCL2, ↑CXCR4, ...all 6 | |
| DNMT3A | | enzyme | | | | 4.94E-02 | ↑CXCR4, ↑EMX1, ...all 4 | |
| 3-methyladenine | | chemical toxicant | | | | 4.95E-02 | ↑BCL2, ↑TP73 ...all 2 | |
| PRKCZ | | kinase | | | | 4.95E-02 | ↓ABCA1, ↑BCL2 ...all 2 | |
| ICMT | | enzyme | | | | 4.95E-02 | ↑CRABP1, ↑GJA1 ...all 2 | |
| ZBTB17 | | transcription regulator | | | | 4.95E-02 | ↑BCL2, ↑NGFR ...all 2 | |
| S100A8 | | other | Inhibited | -2.000 | | 5.00E-02 | ↑BCL2, ↑BMF, ↑... ...all 5 | |
| TP53 | | transcription regulator | Activated | 2.010 | | 5.40E-02 | ↓ASL, ↑BCL2, ↑... ...all 28 | |
| E. coli serotype 0127B8 li | | chemical - endogenous... | Inhibited | -2.219 | | 5.82E-02 | ↑ADRA2A, ↑CH... ...all 5 | |
| TNF | | cytokine | Activated | 2.033 | | 6.33E-02 | ↓ABCA1, ↑ACK... ...all 27 | |
| SIRT1 | | transcription regulator | Inhibited | -2.236 | | 1.19E-01 | ↓ABCA1, ↓CRH, ↑ ...all 7 | |



©2000-2018 QIAGEN. All rights reserved.

| Upstream Regul... | Expr Fold Change | Molecule Type | Predicted Activatio... | Activation z-score | Flags | p-value of ove... | Target molecule... | Mechanistic Netwo... |
|---|---|---|---|---|---|---|---|---|
| SIRT1 | | transcription regulator | Inhibited | [illegible] | | [illegible] | ABCA1, CRH, ... ...all 7 | |
| S100A9 | | other | Inhibited | -2.000 | | 1.22E-01 | BMF, HEY2, ... ...all 4 | |
| prostaglandin E2 | | chemical - endogenous... | Activated | 2.384 | | 1.38E-01 | BCL2, CXCR4, ...all 6 | |
| Insulin | | group | Activated | 2.578 | | 1.54E-01 | BCL2, CRH, ... ...all 7 | |
| PPARD | | ligand-dependent nucl... | | 1.982 | | 2.25E-01 | BCL2, IFI44L, ...all 4 | |
| cyclosporin A | | biologic drug | Inhibited | -2.449 | | 3.77E-01 | BCL2, CXCR4, ...all 6 | |
| RB1 | | transcription regulator | | 1.964 | | 4.40E-01 | BCL2, EOMES, ...all 5 | |
| PTEN | | phosphatase | | -1.972 | | 1.00E00 | BCL2, CXCR4, ...all 5 | |



©2000-2018 QIAGEN. All rights reserved.
